# A Complete Spatial Map of Mouse Retinal Ganglion Cells Reveals Density and Gene Expression Specializations

**DOI:** 10.1101/2025.02.10.637538

**Authors:** Samuel A. Budoff, Alon Poleg-Polsky

## Abstract

Retinal ganglion cells (RGCs) transmit visual information from the eye to the brain. In mice, several RGC subtypes show nonuniform spatial distributions, potentially mediating specific visual functions. However, the full extent of RGC specialization remains unknown. Here, we used en-face cryosectioning, spatial transcriptomics, and machine learning to map the spatial distribution of all RGC subtypes identified in previous single-cell studies. While two-thirds of RGC subtypes were evenly distributed, others showed strong biases toward ventral or dorso-temporal regions associated with sky vision and the area retinae temporalis (ART), the predicted homolog of the area centralis. Additionally, we observed unexpected spatial variation in gene expression within several subtypes along the dorso-ventral axis or within vs. outside the ART, independent of RGC density profiles. Finally, we found limited correlations between the gene profiles of the ART and the primate macula, suggesting divergent specialization between the mouse and primate central vision.

## Introduction

More than a hundred years ago, neuroanatomists described the basic information flow of the retina, conceptualizing light detection in the outer retina by photoreceptors^1,2^, and sequential propagation of information through intermediate layers to retinal ganglion cells (RGCs), which project to the brain^3^. In addition to the differential specialization of its layers along the ‘vertical’ axis, research going back to Thomas Young demonstrated that the *en face* or ‘horizontal’ aspect of the human retina is inhomogeneous, where high-resolution vision in the central retina is supported by the macula and its center, the fovea, which are anatomically and functionally distinct from the peripheral retina^4,5^.

The classic definition of these and other retinal specializations requires locally increased photoreceptor and/or RGC packing and/or differential retinal thickness^6^. Arguably the best-known specialization is found in species with frontal-facing eyes, termed the ‘*area centralis*’. This is a small region of the retina with high RGC density which experiences minimal optic flow during locomotion and is used for binocular fixation and depth discrimination^6–11^. For animals with lateralized eyes, minimal optic flow occurs on the part of the retina that samples from frontal visual fields, which will generally be on the temporal, or for mice, the dorso-temporal retina^6^. For these species, the “Terrain Hypothesis of Vision”, which was developed based on insights from histology, optics, analysis of lifestyle, ecological niche, and eye height from the ground predicts the retinal specialization termed *area retinae temporalis* (ART)^6^.^7–9,12–17^. Relevantly to the mouse, the terrain hypothesis further predicts the absence of lateral streak, a structure common to taller species that regularly view an unobstructed horizon. In addition to gross anatomical specializations, the retina is known to vary in its molecular or subtype spatial distributions^15,17–31^. In accordance with efficient coding principles, s-Opsin expression and enhanced contrast sensitivity in the laboratory mouse photoreceptors clearly demark the ventral retinal domain that samples from the sky^19–23,32,33^. Some RGCs are sensitive to this photoreceptor topography, which influences their center-surround receptive field organization and response properties^34,35^ .

The murine retina doesn’t contain areas with significant differences in thickness and only slight differences in total RGC counts^24,36,37^. A slight *area centralis* that does not relate to optic flow^8,16^ exists just ventral to the optic nerve head^15,24^. Although the most numerous functional RGC subtype, termed W3, is more common in the *area centralis*^24^, a more nuanced examination reveals not all RGC subtypes cover the retina uniformly or favor this central region. Most notably, the alpha sustained (*α*S) RGCs cluster in the dorso-temporal retina^15^, are genetic and functional orthologs of primate midget RGCs - which are highly enriched in the macula/fovea - leading to previous studies proposing the ART to be the rodent homolog of the primate central retina^12,15,16^.

Despite decades of intense research, the field managed to map the spatial distribution of only 17 of the mouse RGC subtypes^15,16,24,26–30,38,39^, out of ∼45 distinct RGC subtypes in the mouse retina identified by single-cell RNA sequencing (scRNAseq)^40–42,41–45^. A limited number of unique genetic markers for RGCs challenges traditional mapping approaches that rely on immunohistochemistry (IHC) and Cre-line labeling. These approaches are further complicated by non-specific labeling, requiring the researchers to resort to labor-intensive approaches like analyzing morphological or functional differences to distinguish between RGC subtypes in a single project^29,46,47^. As an alternative, some studies have used scRNAseq on small retinal regions, successfully differentiating RGC subtype distributions between the primate fovea and periphery, suggesting glimpses into RGC topographies in the mouse^31,45,48–50^. In a more granular approach, two recent tour-de-force studies sequenced individual cells via PatchSeq, revealing functional correlates of some RGC subtypes found in genetic studies^47,51^. Unfortunately, these methods lack the spatial resolution or the throughput to comprehensively characterize RGC coverage across the retina.

Recent advances in high-plex spatial transcriptomics at sub-cellular resolution offer a promising path forward toward mapping all cells in a single tissue^52,53^. A recent demonstration of these techniques in cross-sections of the mouse retina successfully identified all major retinal cell types, but did not clearly characterize their spatial distributions^31^. Here, we developed a novel *en face* cryosectioning technique to preserve spatial distribution patterns^54^ combined with a novel machine learning method for dimensionality reduction to select a high-quality gene panel. This enabled subtype inference from puncta counts measured with 10X Genomics Xenium^53^. Using this approach, we were able to uncover the spatial maps of all 45 RGC subtypes identified in single-cell studies^41^ . Extending rigorous, unbiased geospatial methods with machine learning, revealed four interpretable RGC subtype groupings with distinct retinal coverages. Specifically, about one-third of subtypes preferentially sample either from the ART, or from a previously unrecognized, sky-facing, horizon-avoiding specialization^30^.

Finally, molecular maps showed that in addition to different distributions, these groups varied in their genetic profiles and we identified several non-marker, spatially variable, differentially expressed genes (DEGs) within RGC subtypes. We further focused on five gene families related to synaptic transmission and membrane excitability, comparing their spatial distributions between mice and primates. We found low correspondence in gene expression between mouse ART and anatomical area centralis to primate macula. Although the expression of the voltage-gated sodium channel gene family showed a conserved positive correlation between central and peripheral vision across species^43,45,49^, while GABA receptor (GABAR) expression was anticorrelated. These findings provide detailed subtype and molecular maps of the mouse retina, revealing spatial specialization zones and their potential relevance to human vision.

## Results

### Spatial Transcriptomic Mapping of All 45 RGCs

The retina contains several distinct classes of cells, all of which must be accurately distinguished to enable reliable subtype inference (Figure 1a). To classify RGCs based on expression profiles with the initial 300 genes available on 10X Genomics Xenium, we employed the GraSP algorithm for dimensionality reduction. This algorithm uses neural network ensembles, each trained on datasets containing 50% target and 50% distractor cells (Figure 1b)^53^. We integrated published single-cell datasets^41,43,44^ and ranked genes by their importance in classifying all 130 retinal subtypes. This procedure yielded a list of 225 genes that preserved class and subtype separability, which were supplemented with 75 additional genes of interest encoding synaptic proteins and voltage-gated ion channels.

**Figure 1.**
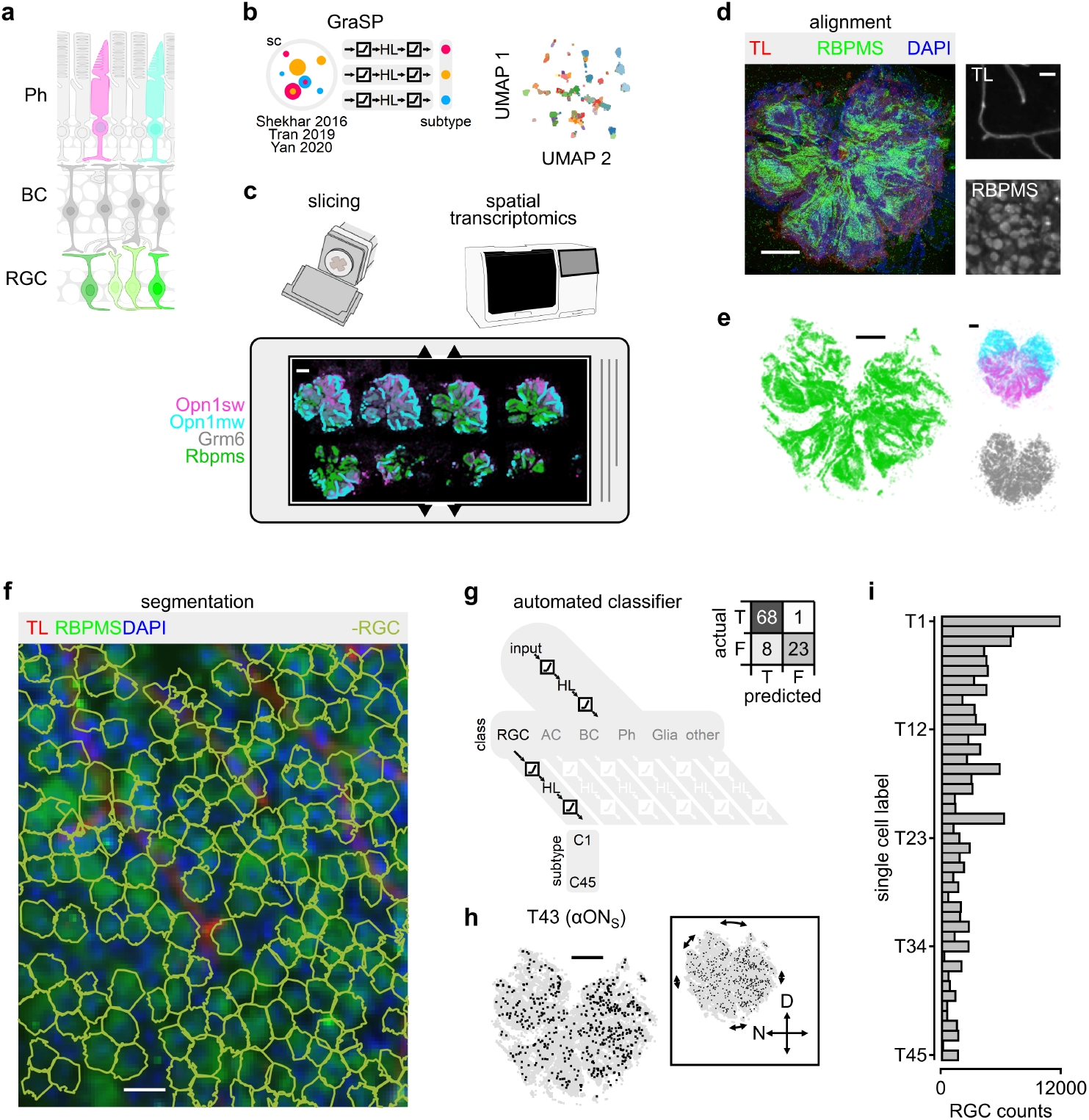
The approach for studying the spatial distribution of RGC subtypes in the retina. **a**. Schematic of retinal organization. Ph, photoreceptors: S-cones (magenta), M-cones (teal), rods (light grey). BC, bipolar cells (dark grey). RGC, Retinal Ganglion Cells (shades of green). **b**. Left, schematic of the Gradient Selected Predictors (GraSP) algorithm, used to identify a set of <300 genes that optimally define retinal subtypes. Right, UMAP projection of RGC subtypes clustered using GraSP-derived genes from the single-cell dataset. Additional panel validation in (Figure S3) and expanded UMAP in. **c**. Representative 10x Genomics Xenium slide showing 8 horizontal slices from one retina, color-coded by cone (magenta/teal), bipolar (grey) and RGC (green) class marker genes. Scale bar – 1mm. **d**. Slices from the same retina as in **c**, stained with Tomato Lectin (TL, blood vessel marker), RBPMS, and DAPI. Images are aligned to the same retinal coordinates. Scale bar – 1 mm. Insets, zoomed-in region showing RBPMS and TL alone, scale bar – 30 *µ*m. **e**. Spatial distribution of genes highlighted in **c**, within the aligned coordinates. Scale as in **d. f-g**. Two-step workflow of ‘CuttleNet’ automated classifier for class and subtype identification of RGCs. **f**. Same histology staining as in **d**, with Baysor-predicted segmentation boundaries of cells identified as RGCs (green). **g**. CuttleNet automated classifier identifies RGC subtypes after initial class identification to distinguish non-RGC cells while avoiding training data inconsistencies between scRNAseq datasets. Inset, confusion matrix showing the accuracy of machine-learning predictions compared to manual RGC identification from RBPMS staining. **h**. Spatial distribution of a single RGC subtype (black) overlayed on all RGCs (grey) in the retina shown in **d-e**. Scale bar – 1 mm. Inset, the data corrected for relief cuts. **i**. Total number of cells identified for each RGC subtype

Retinas from 5 adult C57BL/6J mice (3 females and 2 males) were sectioned into 20 *µ*m thick horizontal (*en face*) slices and mounted on 10X Genomics Xenium slices (Figure 1c) ^54^. Following transcriptomic profiling, the tissue was stained with anti-RBPMS antibody and tomato lectin (TL) to label RGCs and vasculature, respectively (Figure 1d, Figure S1). See supplemental data for RGC-vasculature spatial relationship analysis (Figure S1). IHC images from each tissue slice were registered to the Xenium slide and manually oriented by dorsal-ventral gradients of the *Opn1sw* and *Opn1mw* opsin genes (Figure 1e,f, Figure S2a)^19–23^.

Baysor segmentation^55^ assigned transcripts to cells using nuclei priors (Figure 1f). To differentiate between RGCs and other retinal cells we employed a supervised hierarchical deep neural network for automated classification, CuttleNet (Figure 1g)^53^. CuttleNet consists of a ‘head’ module that identifies cell classes (e.g., Ph, BC, RGC) and dynamically routes them to the corresponding ‘tentacle’ subnetwork to predict specific subtype identities (Figure 1g, Figure S3, S4)^53^. The two-stage inference was important, as the amalgamated scRNAseq training data did not contain identical gene sets between studies^41,43,44^. The performance of Baysor and CuttleNet’s ‘head’ was assessed first by examining relative inferred RGC subtype counts compared to scRNAseq dataset. While enrichment methods applied in scRNAseq studies complicate the estimation of the relative abundance of RGC subtype proportions, we found a good correlation (*R*^2^ = 0.71) between our subtype counts and Tran’s dataset (Figure 1i, Figure S5, S6)^41^, see also^31,40^.

For validation of CuttleNet’s ‘tentacles’, we compared puncta counts with normalized gene expression values of single-cell datasets. Using z-scored gene expression vectors of all 300 genes for each cell class in each data source^41,43,44^ (Figure S3) we computed pairwise distance vectors (Figure S3) that revealed clear separation between cell classes and RGC subtypes (Figure S3). Next, we directly examined RGC inference accuracy by comparing binary class predictions with manual classification based on RBPMS IHC, demonstrating a precision of 89.5% and a recall of 98.5% (Figure 1g, inset, n=255 RGCs). This approach also allowed us to verify Baysor^55^ segmentation, which we found to be more accurate in correctly identifying individual cells in the ganglion cell layer relative to Xenium’s built-in segmentation (Figure S5b,e) and other algorithms^56–63^ (data not shown).

We further examined subtype assignment using genes that nearly exclusively mark specific subtypes: Zic1, expressed in Tran’s subtype cluster 6 (T6)^41^ and Kcnip2, which largely, but not exclusively, marks T45^41,46^ (Figure S5). 96.4% of T6 RGCs classified using CuttleNet contained at least one Zic1 puncta, very similar to the 96.5% T6 cells expressing this gene in Tran’s dataset^41^. Similar analysis showed that Kcnip2 is expressed in 72% of inferred T45 cells, which is lower than 85-100% expression levels found in previous work^41,46^ (Figure S5). This discrepancy could be due to the relative rarity of T45 in the scRNAseq training data compared to T6, suggesting that inferrence accuracy is higher for more common RGC subtypes.

Following RGC subtype classification and slice registration, we projected each retina onto a normalized Cartesian grid centered at the optic nerve head (Figure 1h, Figure S2, Figure S6), allowing us to map the local and global distributions in all 45 genetically-defined RGC subtypes (Figure 2), further characterized below.

**Figure 2.**
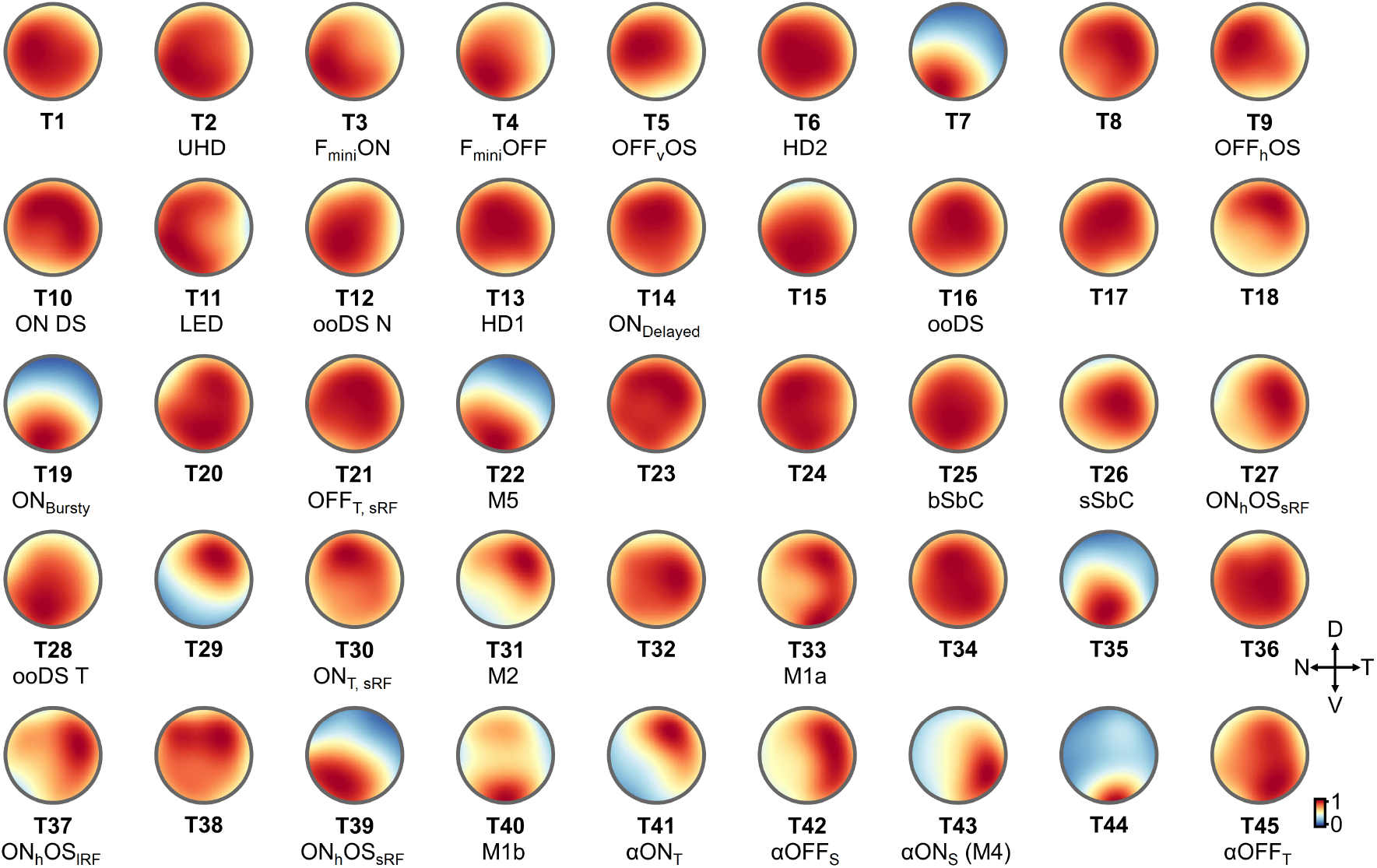
Isodensity maps of all murine retinal ganglion cells. Maps are arranged by genetic clusters proposed in Tran^41^ (in bold). The corresponding functional identity of RGC subtypes from PatchSeq alignment of physiologic subtype identity^47^ to Tran’s cluster scheme is listed below. UHD, Ultra High Definition; OS, Orientation Selective; DS, Direction Selective; oo, ON-OFF; b, bursty; SbC, Suppressed by Contrast; T, transient; S, sustained; sRF/lRF, small/large receptive field.

### Local RGC statistics and mosaics

To assess the local spatial distribution of RGC subtypes, we computed the Voronoi Domain Index (VDRI), the effective radius, and the Nearest Neighbor Regularity Index (NNRI, Figure 3a-d)^14,50,64–68^. We computed the distribution of all subtypes in 14 independent study regions (Figure S7, S8). Statistical significance was tested using bootstrap analysis, where we replaced cells corresponding to a given subtype with the same number of cells randomly sampled from the entire RGC population in the region (Figure S8). Relative to simulating complete spatial randomness, this empirical null distribution incorporates tissue processing artifacts, blood vessels^64,67^, and other hidden variables, such as variable homotypical soma sizes over the retina^15,69^.

**Figure 3.**
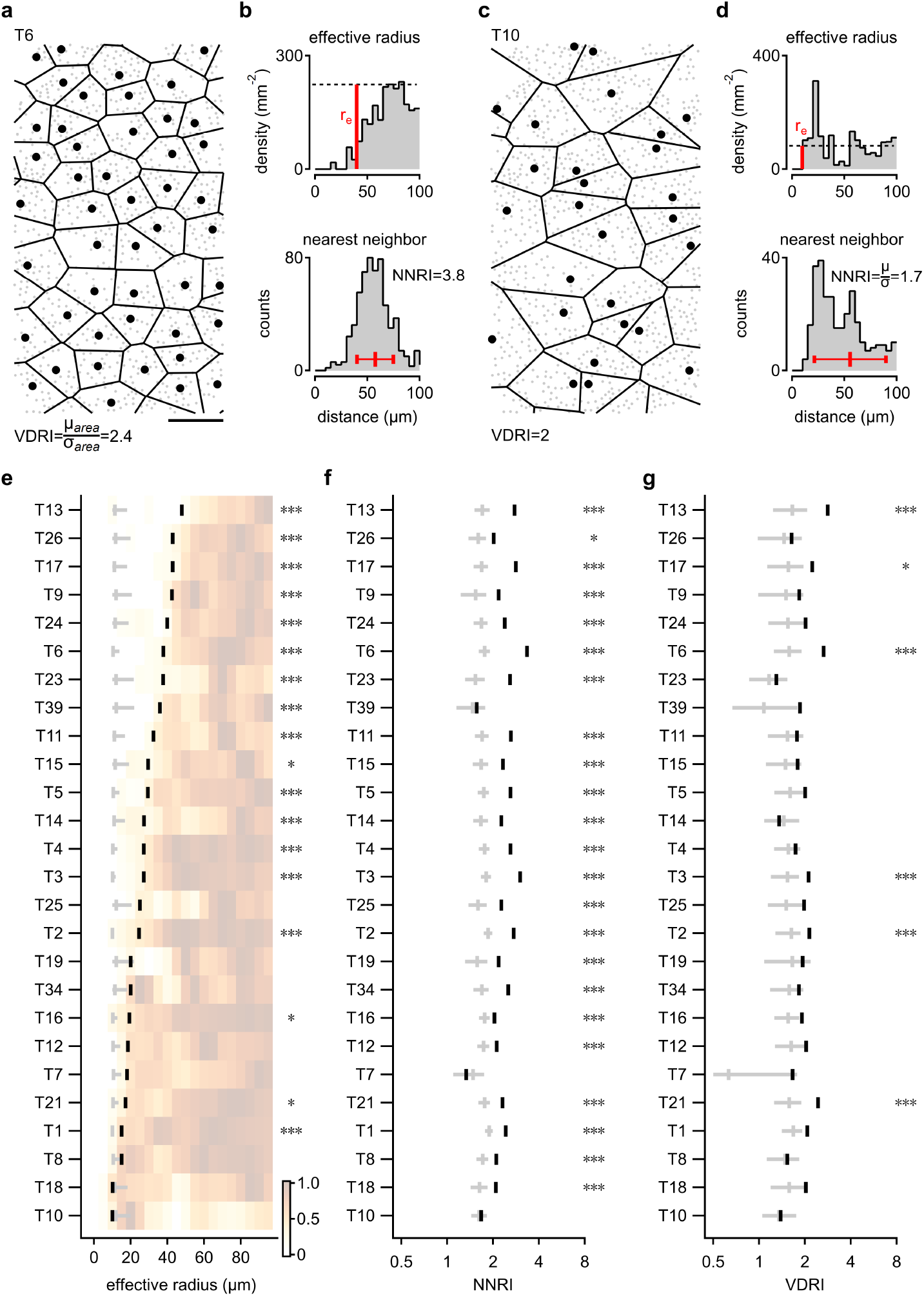
Local statistics of RGC subtypes. **a**. Example Voronoi Domain (VD) analysis for T6 RGCs. VD represents the territory closest to the origin cell than to any other cell; the Voronoi Domain Regularity Index (VDRI) is calculated as the ratio between the mean and standard deviation of all the areas in the study region. Scale bar – 0.1 mm. **b**. The spatial autocorrelation (top) and the distribution of nearest neighbor distances (bottom) for T6, as a function of the distance from the origin cell, used to compute the effective radius (*R*_*e*_) and Nearest Neighbor Regularity Index (NNRI), respectively. **c-d**. as in **a-b**, but for T10, showing less regular spatial distribution. **e**. Effective radius values (black ticks) for RGC subtypes with more than 80 cells in the study regions, overlaid on a heatmap showing normalized spatial autocorrelation. Statistical significance is indicated as follows: ^*^*p* < .05, ^***^*p* < .001, compared to randomly shuffled RGC subtype labels (grey, mean ± 95% CI). **f, g**. Black, observed NNRI (**f**) and VDRI (**g**); Grey, indices computed from bootstrapped populations. See Figure S9 for local statistics of all 45 RGC subtypes.

We observed that 18 RGC subtypes out of the 26 that were well represented in the study regions had effective radii that were significantly larger than predicted by random distributions (Figure 3e, Figure S9). Correspondingly, the NNRI values in the majority of RGC subtypes were higher than expected by chance, although VDRI calculations identified only 6 RGC subtypes that were significantly different from tessellation patterns in random populations (Figure 3f,g, Figure S9). These results suggest that the most common RGC subtypes, for which we could derive a reliable estimation of their local statistics, have an ordered distribution. This information aligns with mosaicism, which is a regular spacing of cells that approximately tile the retina. Interesting exceptions were T7 and T10, which were common in our sample, yet their distribution pattern was statistically random (Figure 3e-g, Figure S9). Segmentation or inference errors could influence local indices, but the evidence for mosaics in many RGCs is a strong indicator of the ability of spatial transcriptomics to identify the local spatial arrangements of most RGC subtypes accurately.

### Geospatial statistics identify subtype specializations in distinct visual fields

The mouse visual system supports a variety of behaviors and functions. For example, predation in mice often involves head and eye movements that position the image of the prey within the binocular zone, which covers approximately 40 degrees of the forward-facing visual fields (Figure 4a,b)^8,16^. As in many other rodents, panoramic visual coverage of the sky is facilitated by lateralized eyes and an enhanced overhead binocular zone, actively compensating for head motion to detect airborne predators (Figure 4a)^6–9^. Previous work linked visual requirements with the spatial distribution of individual RGC subtypes in the mouse. In particular, ipsilaterally-projecting RGCs, including the alpha OFF sustained (*α*OFFS) (T42) and alpha ON sustained (*α*ONS) (T43) densely cover the dorso-temporal retina (Figure 2) and signal prey – with hunting success dramatically reduced with their ablation^8,16^.

**Figure 4.**
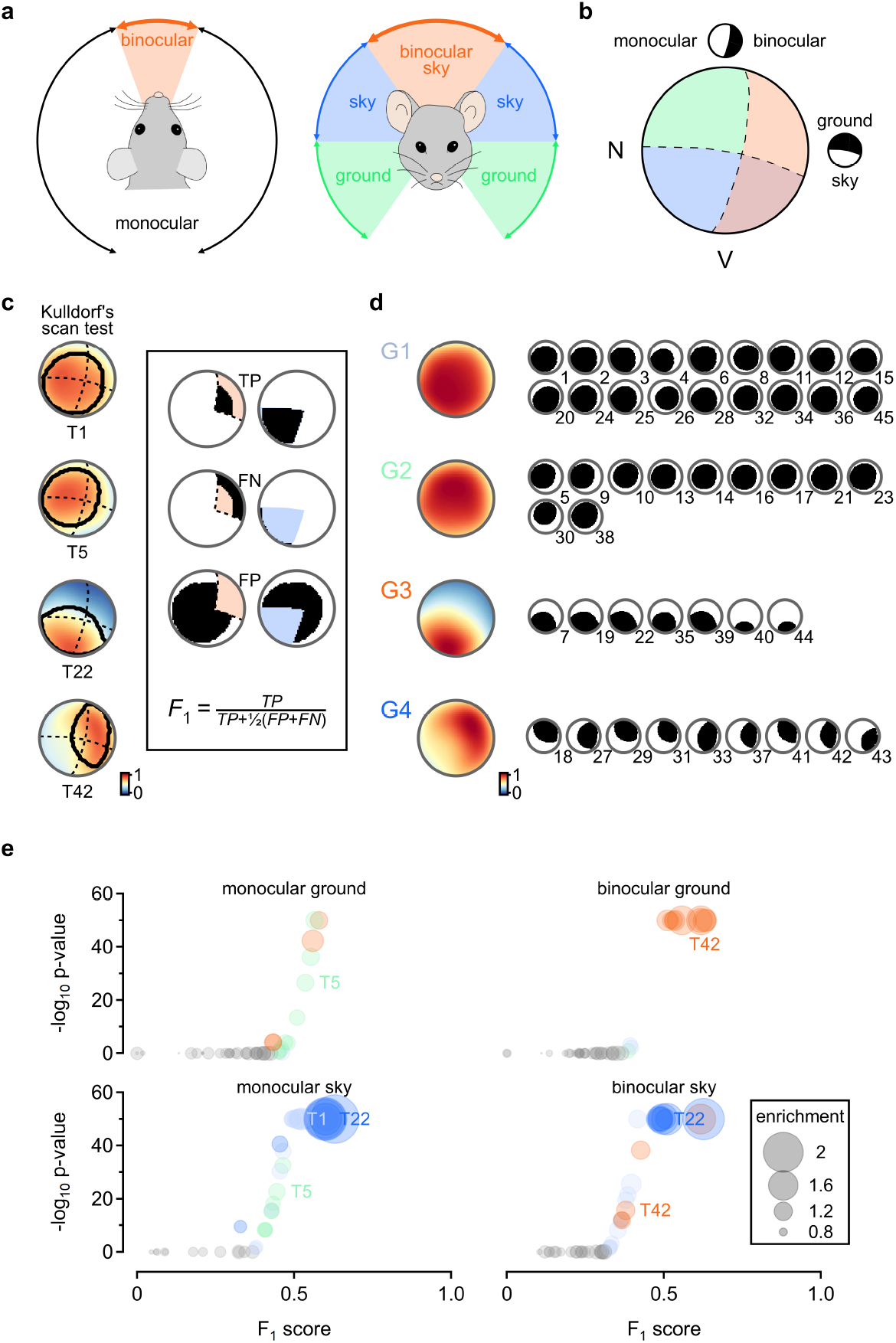
Spatial classification defines regional specializations of RGC subtypes in the mouse retina. **a**. Schematic representation of the mouse’s visual fields along the horizontal (left) and vertical (right) planes, with regions of binocular overlap highlighted in orange. **b**. Projection of the visual fields onto the retina, color-coded as in **a**. Insets illustrate divisions between zones sampling predominantly from the upper (sky) vs. lower (ground) visual fields and monocular vs. binocular regions. Data in **a** and **b** adapted from^8,16^ and shown relative to cells in our study region in Figure S12. **c**. Spatial clustering of example RGC subtypes analyzed using Moran’s I statistic followed by Kulldorff’s spatial scan to identify regions of statistically significantly elevated density (*p* < .001, outlined in black). RGC density maps are as in Figure 2. Dashed curves indicate functional zone boundaries from **b**. Inset, illustration of *F*_1_ score calculation, showing true positive (*TP*), false positive (*FP*) and false negative (*FN*) regions from the overlap of T1 spatial clusters with binocular ground (left) and peripheral sky (right) zones. **d**. Groups of RGC subtypes with differential retinal coverage, identified by clustering F1 scores for each functional zone and RGC subtype. Optimal clustering (*K* = 4) was chosen by an ensemble vote of 9 hierarchical clustering algorithms Figure S9. Left, normalized coverage of retinal space computed from the positions of all RGCs in each group. Right, spatial scan masks for each group member. **e**. Overlap between RGC subtypes and functional retinal zones. Subtypes with significant overlap, determined by Fisher’s Exact test with multiple comparison corrections, are color-coded by group identity as in **f**. Non-significant subtypes are shown in grey. Marker size represents enrichment of RGC coverage, calculated as the ratio of mean density within vs. outside the functional zone.

Given that we have mapped the full RGC population, we next asked whether other regions of the retina are preferentially sampled by specific RGC subtypes. To perform this analysis, we first confirmed that globally, statistically significant spatial clusters (SSSCs) were present in all subtypes, using Moran’s *I*, a test for spatial randomness based on autocorrelation^70^. To locate the positions of the SSSCs, we applied Kulldorff’s scan statistic^71–73^, which evaluates the relative incidence of an observed population within and outside exploratory circles (Figure 4c)^71–73^. We next quantified the relationship between SSSCs and different ethologically relevant subregions of the retina by intersecting SSSC regions with masks representing the binocular sky, binocular ground, peripheral sky, and peripheral ground (Figure S10). For each mask and RGC subtype, we calculated the following metrics: the overlap of the SSSC and the mask (true positive, TP), the part of the mask outside of the SSSC (false negative, FN), the SSSC outside of the mask (false positive, FP), and the part of the study area with neither SSSC nor mask (true negative). The F1 score, the harmonic mean of precision and recall, was then computed to measure the exclusivity of each subtype’s SSSC to a specific visual field (Figure 4c,d).

Unsupervised hierarchical clustering of F1 scores across retinal regions revealed four groups, each composed of RGCs with similar spatial distributions (Figure 4d, Figure S10). Our findings show that the coverage of 65% (29/45) of RGC subtypes extends over most of the rodent retina. These subtypes corresponded to two groups with a somewhat stronger preference either for the region typically occupied by the sky or the ground. Conversely, group 3 consisted of seven RGC subtypes with a clear preference for the ventral retina (thus sampling from the visual sky), including half of the ipRGC types. The other nine subtypes, assigned to group 4, included the other remaining ipRGCs and demonstrated a pronounced preference for binocular visual field in the dorso-temporal retina (Figure 4d,e, Figure S10).

### Spatially DEGs within RGC subtypes are most pronounced on the dorso-ventral axis

RGC characteristics can be shaped by their retinal position^34,35,38,74^. As the spatial RGC groups we identified cover different retinal regions, these groups could vary in gene expression. To explore this question, we chose to focus on the 83 genes encoding proteins involved in synaptic transmission and membrane excitability, only 8 of which were subtype markers selected by GraSP. These gene families were well represented in our panel and convey specific and known functions in neurons. DEG analysis was performed only on genes with mean expression levels > 1 puncta/cell in at least one RGC group (Figure 5a). We found that 6 out of 50 genes that passed the inclusion criteria had significantly different mean expression values (ANOVA, corrected for multiple comparisons), suggesting above-chance genetic similarity between the spatial RGC groups.

**Figure 5.**
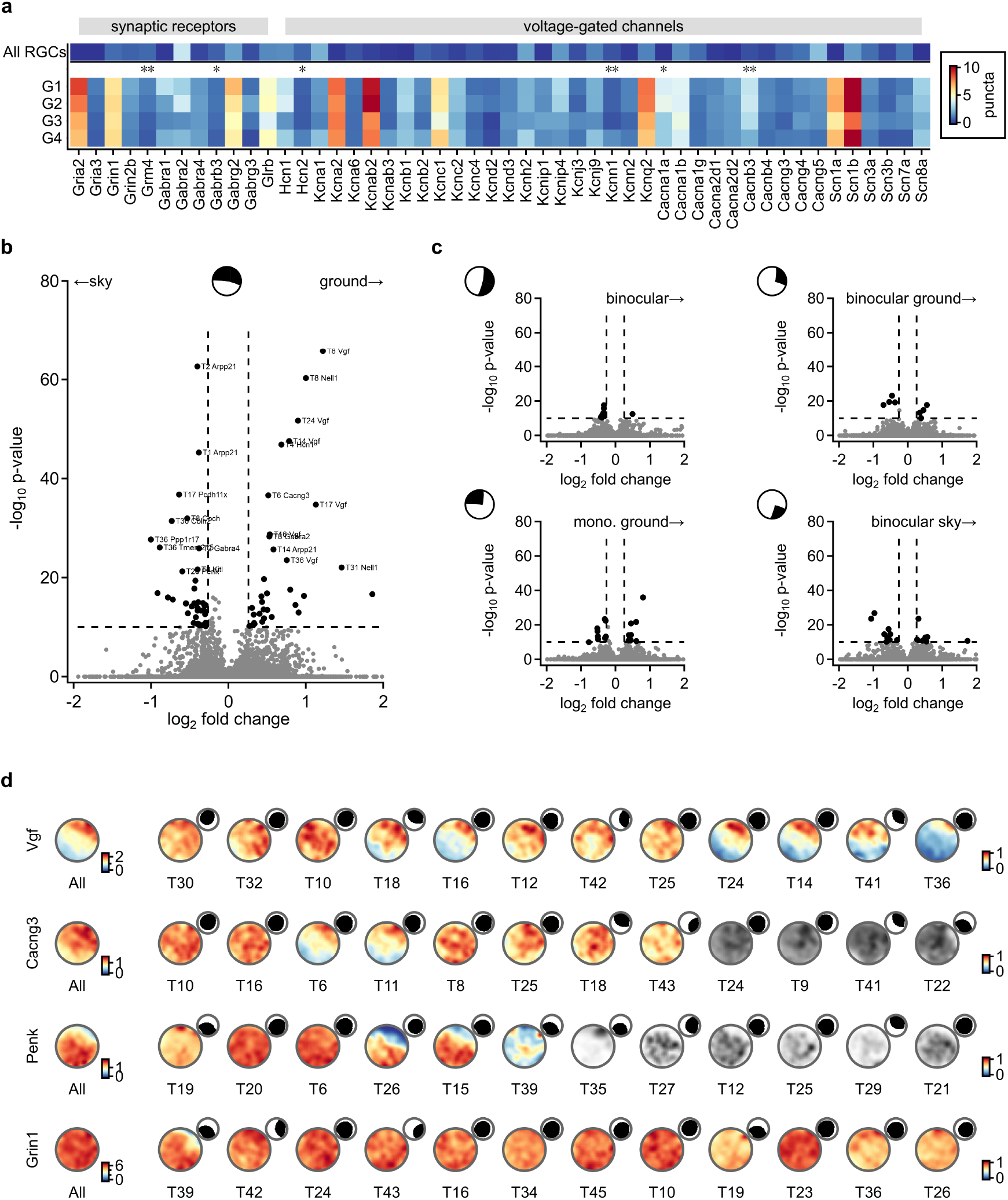
Differential gene expression within RGC subtypes is greatest along the dorso-ventral axis. **a**. Spatially correlated gene expression within SSSC Groups. **b**. Volcano plot illustrating the difference of gene expression across RGC subtypes between the sky and ground sampling portions of the retina (center inset shows the spatial mask). Statistically significant RGC subtype-gene combinations are marked in bold. **c**. as in **b**, for other spatial masks (shown in the upper left). **d**. Example distributions of spatially variable genes (Vgf, Cacng3, and Penk) vs. Grin1, which has similar expression levels across RGCs, for all cells (left) and individual subtypes, sorted by expression levels. RGC subtypes with expression levels less than 1 puncta/cell are color-coded in grayscale. Insets, spatial scan masks of the RGC subtypes.

Next, we focused on individual RGC subtypes and asked if their gene expression varies as a function of retinal topography. To identify DEGs that display within-subtype spatial variability^75^, we pooled RGCs of a given subtype within and outside of different spatial masks (Figure 6a, Figure S11, S12). Most genes in our dataset (260 out of 300) had similar expression across the retina and subtypes (Figure 5b,c). This was anticipated as GraSP-based selection found the most reliable subtype markers. However, a small portion of genes showed statistically different expression levels within at least one RGC subtype (0.9% of all gene/RGC subtype combinations). We found the sky vs ground division was responsible for over half of the DEGs (Figure 5b). Despite the limited size, and engineered bias favoring marker genes of our gene panel, our DEG analysis predicts that several RGC subtypes may have variations in their properties over the retina. Among these are T6 with 7 DEGs and T8, T14, T16, T17 and T36, with 5-6 DEGs.

**Figure 6.**
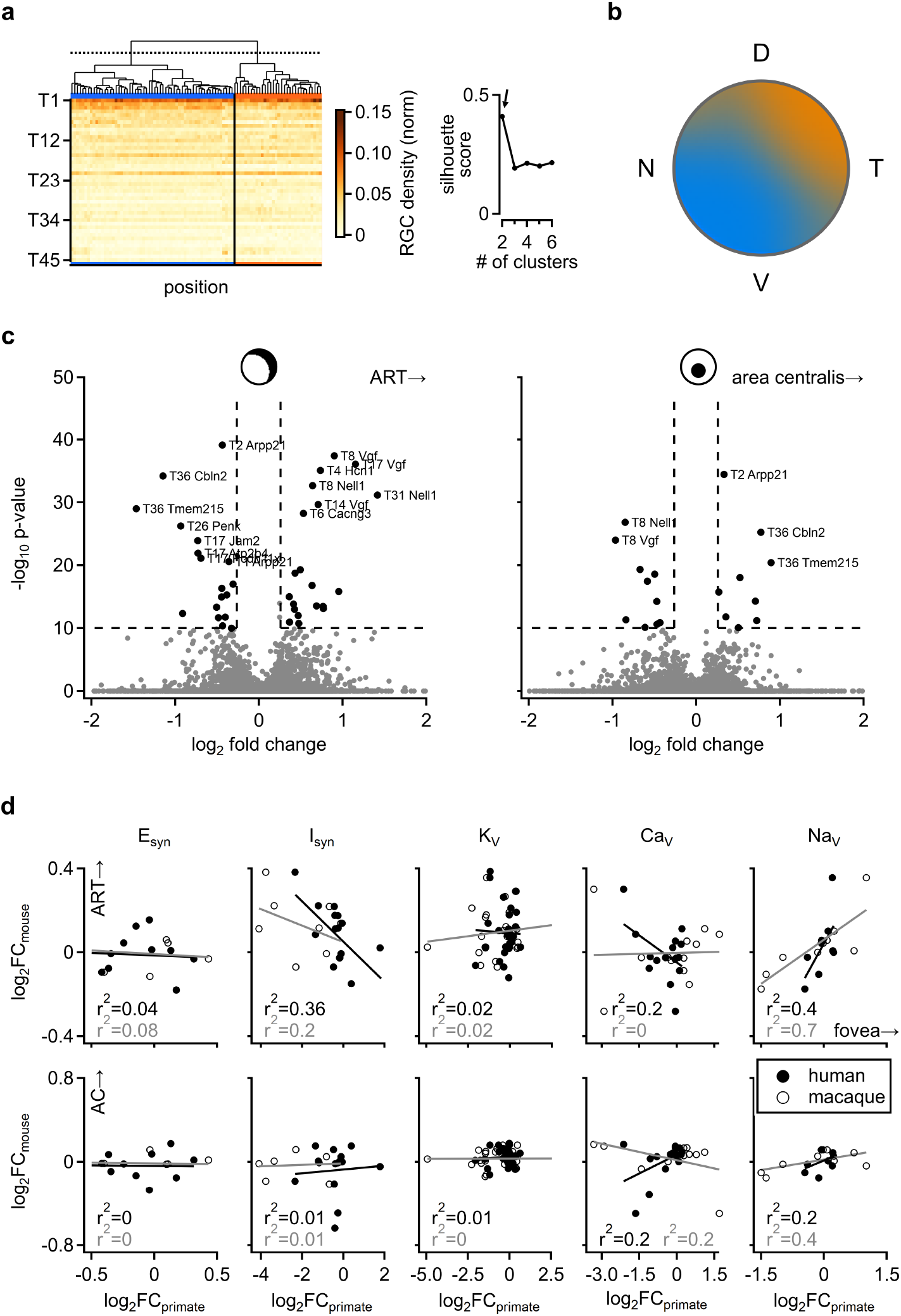
Mouse retina does not have regional homology to primate center vision. **a.** Hierarchical clustering of RGC subtype distribution across the retina, binned into 400 *µ*m × 400 *µ*m squares containing at least 300 RGCs. The optimal number of clusters was determined using silluet score **b**. Cluster assignments from **a**, projected back onto spatial retinal coordinates, smoothed and color-coded by cluster identity. The topography of the dorso-temporal cluster aligns with predicted ART. **c**. Left, DEG analysis of cluster 1 vs cluster 2 (ART) across RGC subtypes. Right, for comparison, an *area centralis* computed from RGC density yields few DEGs. Insets, masks used in the analysis **d**. Correlation between differential expression of genes of interest in the primate and the mouse. Human and macaque data were adapted from published scRNAseq studies from peripheral and central retinal regions. Mouse differential gene expression was computed as in **c**, for ART (top) and *area centralis* (bottom). **25/38**

Several genes demonstrated spatial variation in multiple RGC subtypes, in which case their spatial expression pattern tended to align between subtypes, with most DEG patterns centered in the dorsal-temporal retina (Figure 5d). Rarely, spatially variable genes were found in other retinal locations–for instance, Cosh had higher expression levels in the ventral retina (Figure S11). It is conceivable that this variable gene expression may relate to other observed murine spatial retinal specializations^8,16,34,35^.

### Retinal specializations transcriptomically differ between mice and primates

The mouse is the primary animal model of human eye diseases, including macular degeneration and glaucoma, which affect central and peripheral vision differently^76–78^. This raises the question of how closely the ART —-the proposed evolutionary correlate of the macula in mice—- or the mouse area centralis —-an anatomical correlate of the macula based on higher RGC density and location near the optical axis—-transcriptionally resemble the central retina of primates^6,12^.

To investigate this, we first examined in an unbiased manner whether any part of the mouse retina differs in RGC subtype composition based on subtype densities alone. We first divided the mouse retina into a 400 *µ*m grid and calculated vectors representing the relative densities of RGC subtypes at each spatial location. Next, we clustered these vectors using an ensemble of hierarchical clustering algorithms (Figure S13). This analysis identified two distinct spatial clusters with different RGC subtype compositions (Figure 6a). Interestingly, one of these clusters occupied the dorso-temporal region, closely matching the predicted topographical position of the ART (Figure 6b).

To compare gene expression between the ART-like cluster vs the rest of the retina with that of the primate central vs peripheral retina, we correlated pooled puncta counts from all RGCs in both spatial clusters against gene expression data from primate scRNAseq datasets (Figure 6c,d). As above, we limited the analysis to synaptic proteins and voltage-gated channels that had orthologous genes in the primate datasets. We saw only modest correlations between mouse and primate glutamate receptors and calcium or potassium voltage-gated channels (Figure 6d). However, sodium voltage-gated channel and inhibitory synaptic receptors (GABA and Glycine) subunits significantly correlated and anti-correlated, respectively, between the mouse ART:periphery and primate macula:periphery (Figure 6d). These observations suggest even though the behavioral role the ART and primate macula play is potentially similar, their functional transcriptomic profile is not related.

Next, we repeated the same DEG analysis on the mouse *area centralis*. We found fewer DEGs than from the unbiased ART cluster (Figure 6c). We next regressed the mouse *area centralis* and primate macula, again finding low correlations in functional gene sets. Although, as in the ART, the sodium voltage-gated channels were weakly positively correlated between mice and primates (Figure 6d). Detailed analysis showed that the correlations in the voltage-gated sodium channel gene family were primarily driven by Group 3 subtypes, whose most dense distribution is found far from the ART and area centralis and thus do not reveal specific specializations to these regions (Figure S14).

## Discussion

We used spatial transcriptomics to create a detailed atlas of RGC subtype distributions in the mouse retina. By optimizing *en face* slicing^54^, we preserved large, intact regions of the ganglion cell layer. To address the high cellular and molecular density in our dataset, we implemented a Bayesian segmentation algorithm^55^ and invented novel machine learning approaches for dimensionality reduction and inference^53^ trained on single-cell RNA sequencing^41,43,44^ and fine-tuned with spatial puncta distribution data. This allowed us to identify and map all RGC subtypes in the retina. While the majority of RGCs exhibited relatively uniform spatial distribution, about one-third showed a strong preference for either ventral or dorso-temporal retinal zones. Interestingly, gene expression also varied across the retina but was largely uncorrelated with the spatial profiles of the RGC subtypes expressing those genes. Finally, the topographic distribution of genes of interest expressed in primate central vs peripheral retina was regressed against both the behaviorally homologous *area retinae temporalis* and anatomically homologous *area centralis*. In both cases, we found weak correlations between the genetic profiles of the primate central versus peripheral retina, suggesting divergent specialization between the mouse ART and the primate macula.

We found close agreement between our results and published RGC subtype distributions (Figure 2). In the temporal retina, we reproduced the high densities of *α*ONS, alpha ON transient (*α*ONT) and alpha OFF sustained (*α*OFFS) RGCs, corresponding to T41, T43, and T42 respectively^15,39^. The ventral retina is enriched in several subtypes: W3 cells^24^ genetically correspond to multiple Tran clusters (T2, T3, T4, T6, T21, T23, T30 and potentially T1 and T13)^41,47,51^, as well as FminiOFF (T3), FminiON (T4), and FmidiOFF (T28)^26^. As expected, FmidiON (T38) cells showed a dorsal bias^26^, and J-RGCs (T5) exhibited a central clustering pattern^38^. The intrinsically photosensitive RGCs (ipRGCs) showed distinct regional patterns: M1 ipRGCs (T33, T40) displayed a bimodal distribution with peaks in both ventral and temporal retina, M2 ipRGCs (T31) showed a dorso-temporal peak, M4 ipRGCs (T43) had a centro-temporal peak, and M5 ipRGCs (T22) were strongly concentrated in the ventral retina, all in line with previous work^29,39,79^. Of the ∼17 murine RGC subtypes historically mapped, the only subtype we observe conflicting results with is the *α*OFFT, T45. This subtype was described as uniformly distributed with a slight temporal bias^15^. Instead, we found a modest ventral-temporal peak, which aligns with a proposed physiological role of T45 as a detector of approaching aerial predators^25^. Moreover, the observed RGC subtype proportions in our dataset were more similar to those from scRNAseq studies^40,41^ compared to the previous spatial transcriptomic classification of retinal cross sections^31^. Intriguingly, we recapitulated the increased T16 and T21 prevalence relative to scRNAseq datasets observed by this group. Lastly, the RGC maps generated in this study closely align with the distributions reported by Tsai and colleagues in a recent spatial transcriptomic investigation. Despite substantial differences in techniques and analytical methods, this consistency supports the accuracy of our subtype identification and the topographical organization of various RGC populations^80^.

Efficient coding theory proposes that sensory systems evolve to encode environmental information accurately while minimizing resource consumption^33^. According to this theory, homotypic RGCs are expected to tile the retina without redundant sampling^81^. Consistent with this prediction, numerous studies have demonstrated that retinal cell types are organized in non-random mosaic patterns^15,24,27,28,67,82–88^. However, like all nocturnal animals, mice rely on vision that performs effectively under scotopic conditions. In such environments, redundant visual information becomes an advantage, as the ability to detect and discriminate rare photons can be critical for survival. Increased sensitivity in dim light can be achieved through various adaptations, such as clustering photoreceptors; exemplified by the “macroreceptors” of *Scopelarchus guntheri*^7,89^. Another strategy is to enhance RGC receptive fields via gap junctions or a reduction in surround inhibition^6,7,89–91^, prompting visual ecologists to point out mosaics are not the expectation for nocturnal animals with poor optics^6,7,92^. We note that as some RGC subtypes were not well represented in our study regions, we could not draw conclusions on the local spatial statistics of all RGCs. Yet, our findings match the conclusions reached in studies employing rigorous spatial statistics showing that regular mosaics occur for some, and likely most, RGC subtypes but are not a general organizational principle^64–68^.

What is a subtype? This is a fundamental question not only in the study of the retina, but in neuroscience in general. Given our challenge to the idea that mosaics define subtypes, we wanted to delve deeper into this question. To date, the three complimentary^47^ techniques to define subtypes include morphology^3,69,93,94^, physiology^81,95–97^, and genetic profile^12,41–46,48–50^. Individual RGC subtypes’ morphology^30,38,98,99^ and physiology^34,74^ can vary as a function of cell position in the retina. scRNAseq studies dramatically increased the number of putative RGC subtypes than was appreciated in morphological or functional examinations. Potential “overcounting” of RGC subtypes could be explained by spatially variable genetic profiles of RGCs. Spatially variable gene expression has been reported within the same “subtype” in the retina^12,45,48–50^, and other brain regions^100–107^. In this scenario, a single cellular phenotype could be erroneously divided into multiple subtypes in scRNAseq analysis, which has no access to topographic information. Our examination of RGC coverage and DEG analysis concretely disproves the possibility that scRNAseq overestimated the number of RGC subtypes in the mouse. First, while retinal coverage of several RGC subtypes could form a continuum (for example, subtypes belonging to specialization group 3 with subtypes from group 4), these RGCs either had radically different gene expression patterns or matched characterized distributions of specific RGC populations. Second, in addition to the presence of mosaics in many genetically-defined RGC subtypes, we show that the number and identity of DEGs are not sufficient to drive erroneous subtype assignment.

Unexpectedly, we found that many of the spatially variable genes were expressed differently within a narrow region of the dorso-temporal retina, a zone that had statistically different RGC subtype distributions, closely matching the expected position of the ART^6,8,12,16^. Yet, despite previous proposals linking the ART to the primate central vision based on function and orthologous RGC subtype topography^8,12,15,16^, our findings show that genes encoding synaptic receptors and voltage-gated channels do not show matching differential expression between mice and primates. The only strongly correlated gene family was the voltage-gated sodium channels, and a detailed analysis shows that this difference is driven by a small population of RGCs in the ventral retina.

Looking ahead, this work opens several promising avenues for future research. First, the use of more advanced transcriptomic techniques that do not require *en face* sectioning will eliminate slicing artifacts and allow mapping of larger retinas. Second, our gene panel was designed to classify all retinal cell subtypes, including bipolar and amacrine cells. However, current segmentation methods did not adequately resolve the inner nuclear layer to our satisfaction^56–63^. Improved segmentation would clarify the spatial relationships between photoreceptors, bipolar, amacrine, and ganglion cells within the same tissue, enabling the development of more accurate models of retinal physiology. Such analysis could shed light on how the specializations identified in the RGC layer are organized in relation to upstream circuits. This, in turn, would provide deeper insights into the computational strategies employed across different retinal regions, which is crucial for understanding vision in mice and, eventually, humans.

## Methods

### En Face Retinal Cryosectioning and Molecular Profiling

All animal procedures were conducted in accordance with U.S. National Institutes of Health guidelines, as approved by the University of Colorado Institutional Animal Care and Use Committee (IACUC). Mice were housed at a 12 hour light/12 hour dark cycle room set to fixed ∼22 ° C temperature and 40–60% relative humidity. At the time of terminal surgery, mice were deeply anesthetized with isoflurane. Anesthesia was verified via toe pinch and sacrifice was performed with cervical dislocation followed by decapitation. After sacrifice, the temporal pole of each globe was branded and the left eye was extracted using forceps before being immediately submerged in Ames solution (Sigma-Aldrich, A1420). Retinas from five (3 female + 2 male) C57BL/6 mice were dissected and cryopreserved following the *en face* sectioning protocol detailed in^54^. Briefly, globes were cleaned of muscle, fascia, and the optic nerve before the anterior chamber was removed. Relief cuts were made in the resulting eye cups to allow flattening of the retina with the sclera attached before flash freezing the tissue with the ganglion cell layer pressed flat against a coverslip and the tissue embedded in OCT medium (Fisher Scientific, 23-730-625). Retinas were cryosectioned at -18^°^ C to generate 20 *µ*m sections collected on Xenium slides (Figure 1c) from photoreceptors toward RGCs^54^. Molecular profiling was performed using the 10X Genomics Xenium platform using a custom 300 gene panel in which 225 genes were selected in an unbiased manner by the GraSP algorithm^53^, supplemented with manually selected 75 genes encoding synaptic proteins and voltage gated channels (Figure S1a). Xenium *in situ* hybridization was conducted as per manufacturer protocols by the Human Immune Monitoring Shared Resource at the University of Colorado Anschutz Medical Center. After Xenium processing, IHC with rabbit-anti-RBPMS (Invitrogen, MA5-46928) + goat-anti-rabbit AF549 secondary (Invitrogen, A-11012) to label retinal ganglion cells, and Tomato Lectin (TL) conjugated to AF649 (Invitrogen, L32472) to visualize vasculature was applied to the tissue on the Xenium slides (Figure 1d,f, Figure S2a) before a coverglass was applied using SlowFade™ Glass Soft-set Antifade Mountant (Invitrogen, S36917-5X2ML). IHC images were acquired on a Keyence BZX800 epifluorescent slide scanning microscope with optical sectioning via structured illumination. Raw Keyence images were stitched into full slide z-stacks with custom code implemented in Python. Max projected images were used for all analysis.

### Retinal Ganglion Cell Layer Reconstruction

IHC images of all retinas were printed on transparencies to enable manual registration of all retinal sections. Registration parameters, including rotation angle, tissue flipping, and relative translocation were recorded and used to digitally register all tissue slices in IgorPro (Wavemetrics). Digitally registered retinas were rotated to align the dorsal ventral axis via the s-opsin and m-opsin gradient (Figure 1e, Figure S3a). Nasal-temporal orientation was recorded during tissue preparation. All biological replicates were aligned on a common cartesian plane, centering each biological replicate optic nerve head at (0,0) to allow for normalization (Figure S3b). For global spatial analysis, relief cuts were digitally filled in by radially spreading retinal cell centroids to circularize all retinas to the extent tissue was available (Figure 1h, Figure S3c). For local spatial statistical analysis, all of these transformations are omitted. See supplement for additional registration validation.

### Segmentation and Classification

The Baysor algorithm^55^ was used to predict cell boundaries on the basis of gene expression (Figure 1f), using the Xenium prediction of nuclear overlap as a prior. To enable Baysor segmentation, the Xenium explorer graphical user interface was used to manually annotate ganglion cell layer regions guided by expression of the bipolar cell marker Grm6, the amacrine cell marker Prox1 (which is highly expressed in the inner plexiform layer), the ganglion cell marker Rbpms and the protein stain for RBPMS. Raw xenium data files were subset with a custom python script to extract the puncta measured in these ganglion cell layers. Following segmentation, a custom python script was used to consolidate all gene counts into an expression matrix with each row representing a cell and each column a gene (1434195, 300). The CuttleNet Neural network^53^ was used for inference to predict subtype classification (Figure 1g, Figure S1). CuttleNet was initially trained on the scRNAseq training dataset. Cell class (Glia vs. Photreceptors vs. Bipolar Cells vs Amacrine Cells vs RGC) reliability was assessed in a custom python script in which a human was presented a Baysor segmentation outline aligned with an IHC image. The subject was then prompted to classify the segmentation boundary as containing an RGC based on the RBPMS stain or not. Using these 255 human classifications, a confusion matrix was constructed to quantify CuttleNet’s performance. In a separate analysis to assess how well the segmentation boundaries themselves were, a region of the IHC image stack was manually labeled by a human and compared with Baysor segmented regions. This procedure was also performed with other segmentation algorithms, including the default DAPI-based algorithm employed by Xenium. We next measured the Xenium distribution of puncta levels in RGCs and correlated these against the scRNAseq dataset, providing a scaling factor for each gene. After applying this scaling factor to the scRNAseq dataset, we fine-tuned CuttleNet to improve subtype classification. A custom R script was used to filter the dataset to include only RGCs, and to calculate the homotypical nearest neighbor distances in order to merge cells of the same subtype that were biologically implausibly close together^69^. The final dataset count matrix included 130,575 RGCs. Final quality control validations focused on assessing subtype inference. We first compared puncta counts with normalized gene expression values by computing the z-scored gene expression of all 300 genes in each cell class for both data sources (Figure S1a). Pairwise vector distances between cell classes between datasets were then computed to identify linear trends in support of inference accuracy. Next, we visualized the puncta of Zic1 and Kcnip2 against Baysor segmentations and computed the frequency with which these puncta were in the segmentation regions inferred to be T6 and T45, respectively. This procedure was also performed on the Xenium mask regions for additional validation of Baysor segmentation (Figure S2). Finally, the proportion of each cell subtype predicted in the single-cell dataset and our own was compared to identify how well the different census sources aligned.

### Geospatial Statistical Analysis

#### Data Preparation

Normalizing the 5 biological replicates, following CuttleNet classification, we analyzed the distributions of the 45 genetically-defined subtypes across the surface of the retina. Replicates were binned into 400 *µ*m^2^ regions and the total RGC count in the region (Figure S3d) was used as the normalization factor to produce a per-subtype density. The density maps were then smoothed using a 2d Gaussian kernel (Figure 2).

#### Global Spatial Statistical Clustering

To assess the global spatial clustering of RGC subtypes, we computed Moran’s *I* statistic^70^ for each normalized and smoothed subtype density map. Moran’s *I* is a measure of global spatial autocorrelation that quantifies the degree to which similar values cluster together in space. The statistic is defined as:

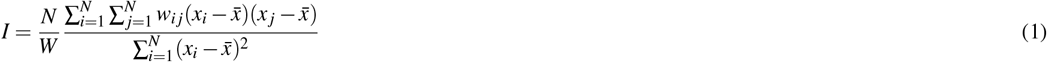

where *N* is the number of spatial units, *x*_*i*_ represents the normalized density value at location *i*, 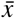 is the mean density across all locations, *w*_*ij*_ are the elements of a spatial weights matrix with zeros on the diagonal, and *W* is the sum of all spatial weights 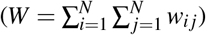.

For each density map, we constructed a queen-contiguity spatial weights matrix, where *w*_*ij*_ = 1 if cells *i* and *j* share either a border or vertex, and *w*_*ij*_ = 0 otherwise. This approach relates to Tobler’s first law of geography^108^, which states that nearby observations are more related than distant ones. The resulting weights matrix was row-standardized to ensure each row sums to unity. Statistical significance was assessed through Monte Carlo simulations (*n* = 999). All of these calculations were performed using the `moran.test` and `moran.mc` functions from the `spdep` R package. For each density map, we tested against the null hypothesis of no spatial autocorrelation (*I* ≈ 0) with the alternative hypothesis of positive spatial autocorrelation (*I* > 0). The expected value of Moran’s *I* under the null hypothesis is:

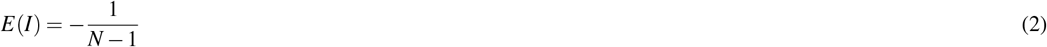

Values of *I* significantly greater than *E*(*I*) indicate positive spatial autocorrelation (clustering), while values significantly less than *E*(*I*) indicate negative spatial autocorrelation (dispersion). We considered the results significant *p* < 0.05 after controlling for multiple comparisons using the Benjamini-Hochberg procedure.

#### Defining The Extent of Spatial Statistical Clusters

To detect localized clusters and define their spatial organization, we employed Kulldorff’s spatial scan statistic^71,72^ implemented through the `SMERC` R package. This method systematically evaluates circular windows of varying sizes across the retinal space to identify regions of significantly elevated cell density. For our implementation, we scaled the normalized grid values by a factor of 10,000 to approximate count data suitable for the scan statistic. We did not use our raw count data as this would not account for our biological replicates.

Let *G* denote the geographic space of our normalized maps. *Z* is a scanning windows *Z* ⊂ G. *µ*(Z) represent the window’s population. For a given subtype, a cell can either be considered inside the scan widow, or outside it, with the respective probabilities for these states being *p* and *q*. The random number of cells in the set *A* ⊂ G is N(A) ∼*P*_*o*_(*P*_*µ*_ (*A* ∩ Z) + q_*µ*_ (*Z* ∩ Z^C^)) ∀A under the inhomogeneous Poisson model. The null hypothesis for this model is thus *H*_0_ : *p* = *q* while the alternative is *H*_1_ : *p* > *q, Z* ∈ ℤ. The choice of using the Poisson model instead of the Bernoulli model relates to our assumption that cell types are distributed in a continuous manner over the surface of the retina, as comes from the expectation of mosaicism. Under the Poisson model, for each candidate window, we calculated a likelihood ratio statistic:

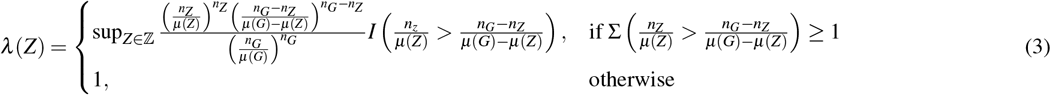

where:

- *n*_*Z*_ is the observed number of cases within window *Z*
- *n*_*G*_ is the total number of cases across the study region
- *µ*(*G*) is the total population of the study region
- *µ*(*Z*) is the total population in the scan window
- *I*(·) is the indicator function

The scanning process was implemented with the following parameters:

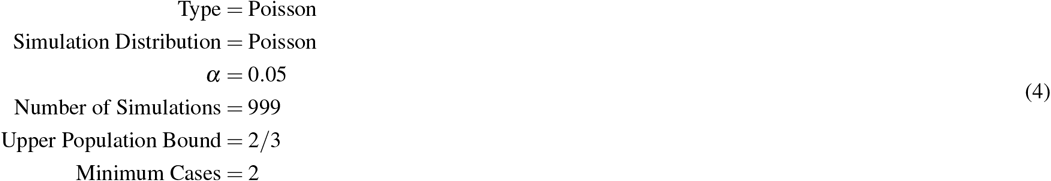

Statistical significance was assessed through Monte Carlo hypothesis testing, where the null hypothesis posits that the observed cases follow an inhomogeneous Poisson process with intensity proportional to the underlying population density. For each window *Z*, we computed:

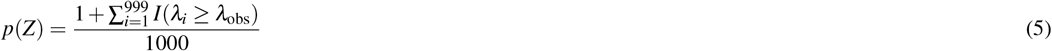

where *λ*_*i*_ represents the maximum likelihood ratio for the *i*th simulated dataset, and *λ*_obs_ is the observed maximum likelihood ratio. Clusters were considered significant at *p* ≤ 0.05. Secondary clusters were identified iteratively, with non-overlapping significant clusters ranked by their likelihood ratio statistics.

#### Extending Geospatial Statistics with Biologic Meaning

To relate the spatial organization of RGC subtypes to ethologically relevant regions of the mouse visual field, we defined a set of masks representing the binocular zone, visual horizon, and peripheral visual field based on prior studies. The ipsilateral projection zone mask was constructed to match the area (∼22% of the visual field) and shape reported by^16^ based on anterograde tracing. The binocular, and visual ground and visual floor zones were assessed based on^8^. The peripheral visual field was defined as the complement of the binocular zone, while the visual sky the complement of the visual ground (Figure S12).

For each RGC cluster *C* and visual field mask *M*, we quantified their spatial association using precision (*P*), recall (*R*), and F1 score (*F*_1_) metrics implemented in base `R`. Given a binary cluster mask and visual field mask, these metrics are defined as:

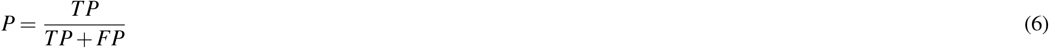

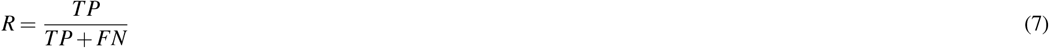

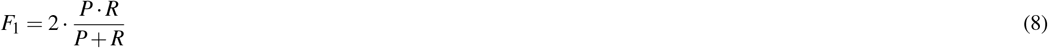

where *TP* (true positives) represent pixels present in both cluster and mask, *FP* (false positives) represent pixels present in the cluster but not the mask, and *FN* (false negatives) represent pixels present in the mask but not the cluster.

To assess the statistical significance of spatial associations, we employed Fisher’s exact test using the `stats` package in `R` on the resulting 2×2 contingency tables. For each cluster-mask pair, the contingency table takes the form:

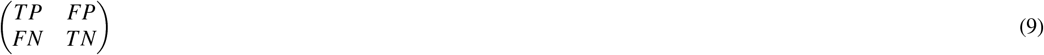

Fisher’s exact test calculates the probability of observing this particular arrangement under the null hypothesis of independence between cluster and mask assignments. The probability is computed using the hypergeometric distribution:

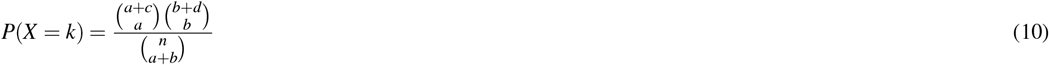

where *n* = *a* + *b* + *c* + *d* is the total number of pixels in the valid study region, and:

- *a* represents true positives (TP)
- *b* represents false positives (FP)
- *c* represents false negatives (FN)
- *d* represents true negatives (TN)

The test also yields an odds ratio (*OR*), which quantifies the strength of association:

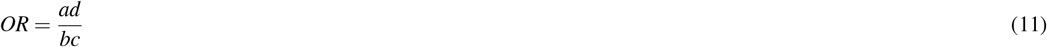

For our analysis, we used a one-sided alternative hypothesis (*H*_*a*_ : *OR* > 1) to identify clusters significantly enriched in specific visual field regions. The resulting *p*-values were adjusted for multiple comparisons using the Benjamini-Hochberg procedure with a false discovery rate of 0.05.

#### Unbiased Clustering of Specializations

The resulting F1 scores were analyzed using hierarchical clustering implemented through the `cluster` and `dendextend` packages in `R`. For a set of *n* subtypes, we computed pairwise distances *d*_*i j*_ between subtypes *i* and *j* using three distance metrics from the `cluster` package:

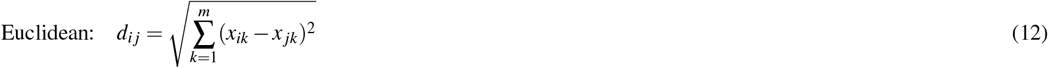

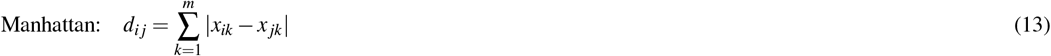

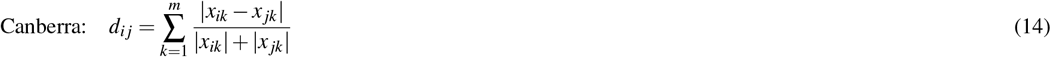

where *x*_*ik*_ represents the F1 score of subtype *i* with mask *k*, and *m* is the number of masks.

For hierarchical clustering, we employed three distinct linkage methods using the `hclust` function in `R` to construct the dendrogram. Each method defines how the distance between clusters is computed during the agglomerative process:

`ward.D` and `ward.D2` minimize the total within-cluster variance. For two clusters *r* and *s*, the linkage distance *d*(*r, s*) is given by:

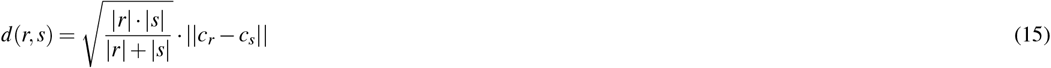

where |*r*| and |*s*| are the sizes of clusters *r* and *s* respectively, *c*_*r*_ and *c*_*s*_ are ||·||their centroids, and denotes the Euclidean norm. `ward.D2` applies a square root transformation to the dissimilarities before clustering.

Complete linkage (maximum linkage) defines the distance between clusters as the maximum distance between any pair of observations:

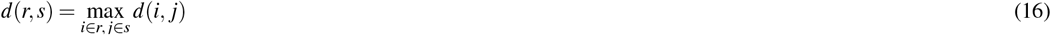

The final clustering solution was selected based on the silhouette score computed using the `cluster` package. For an observation *i*, let *a*(*i*) represent the average dissimilarity between *i* and all other points in its cluster *A*:

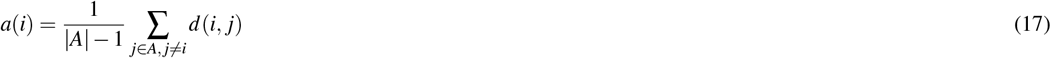

For each other cluster *C*, let 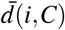 be the average dissimilarity of *i* to all observations in *C*:

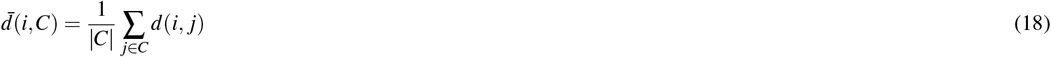

The silhouette coefficient *s*(*i*) is then computed as:

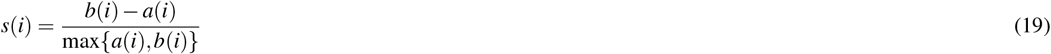

where *b*(*i*) is the minimum average dissimilarity to any other cluster.

The optimal number of clusters was determined by the mode of the maximal silhouette scores across all method combinations (Figure S10, S14), providing a robust consensus clustering solution. Clusters were considered significantly enriched in a visual field region when they exhibited both statistical significance (adjusted *p*-value < 0.05) and practical significance (odds ratio > 1.5).

### Local Spatial Statistics

To analyze the spatial organization of RGC subtypes, we first identified high-quality regions in the retinal slices using Xenium Explorer. Selection criteria focused on areas that were flat, rich in RGCs, and free from tissue deformities, tears, or other visible artifacts. These regions were predominantly located near, but not including, the optic nerve head. A total of 14 distinct regions (Figure S7) were manually annotated using the lasso tool and included in subsequent analysis. Unlike the preceding analysis of global statistics, which relied upon normative, smoothed maps combining all 5 biological replicates and all slices from a given replicate, each region was treated as an independent observation window *ω*.

For each cell type, we computed the Density Recovery Profile (DRP) following Rodieck’s method. The DRP characterizes the spatial organization of cells by analyzing the density of cells as a function of distance from reference cells (Figure S15. For a given reference cell type *r* and test cell type *t*, we:

1. Divided the space around each reference cell into concentric annuli of width Δ*r*
2. For the *i*th annulus, calculated:

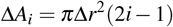

where Δ*A*_*i*_ is the area of the *i*th annulus
3. Computed the density in each annulus:

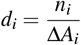

where *n*_*i*_ is the count of test cells in annulus *i*
4. Calculated the expected count in each annulus:

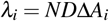

where *N* is the total number of cells and *D* is the mean density
5. The effective radius (*R*_eff_) was determined as the distance at which the observed density first recovers to the mean density after dropping below it:

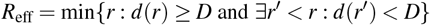 To assess the statistical significance of spatial relationships, we implemented a bootstrap analysis with *n* = 100 iterations. For each cell type pair, we:
6. Calculated the observed ratio:

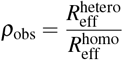

where 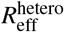 is the effective radius for heterotypic interactions and 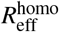 for homotypic interactions
7. Generated bootstrap distributions by resampling distances with replacement
8. Computed confidence intervals using the 2.5th and 97.5th percentiles of the bootstrap distribution
9. Calculated two-sided p-values:

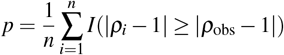

where *ρ*_*i*_ is the ratio from the *i*th bootstrap sample
10. Applied FDR correction using the Benjamini-Hochberg procedure for multiple comparisons

For each RGC subtype within annotated regions, we quantified spatial organization using two more complementary metrics: the Nearest Neighbor Regularity Index (NNRI) and the Voronoi Diagram Regularity Index (VDRI). The NNRI assesses local spatial patterns by computing:

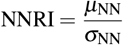

where *µ*_NN_ is the mean nearest neighbor distance and *σ*_NN_ is the standard deviation of nearest neighbor distances between cells of the same subtype. This was implemented using the `spatstat` package’s `nndist` function with edge correction (Figure S8):

The nearest neighbor distance for each cell is calculated as:

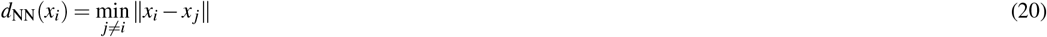

where *x*_*i*_ represents the coordinates of cell *i*.

The VDRI evaluates spatial territory organization through:

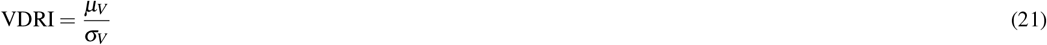

where *µ*_*V*_ is the mean Voronoi polygon area and *σ*_*V*_ is the standard deviation of Voronoi polygon areas. To ensure reliable territory measurements, we implemented a buffer zone exclusion (Figure S8):

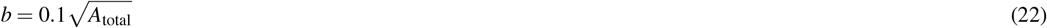

where *b* is the buffer width and *A*_total_ is the total region area. Cells within distance *b* of the region boundary were excluded from VDRI calculations. For statistical validation, we implemented a Monte Carlo bootstrap approach with the following steps:

Statistical significance was evaluated through a Monte Carlo bootstrap approach:

1. For each subtype in region *ω*, generate *n* = 100 random spatial arrangements:

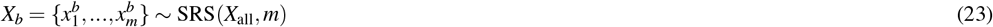

where *X*_*b*_ is the *b*th bootstrap sample, *m* is the observed subtype population size, and SRS denotes sampling without replacement from all cells *X*_all_.
2. For each bootstrap sample, compute NNRI and VDRI:

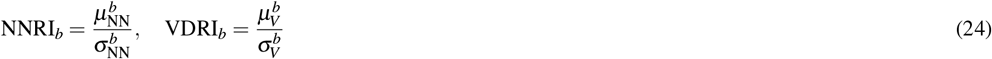
3. Calculate empirical p-values:

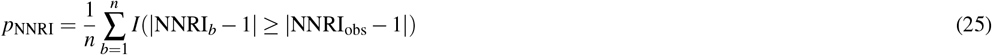

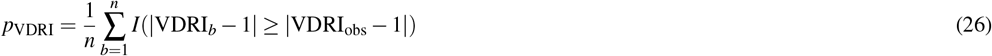
4. Compute 95% confidence intervals:

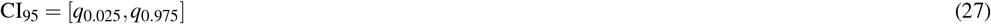

where *q*_*α*_ represents the *α*th quantile of the bootstrap distribution.

Statistical significance was assessed through paired t-tests comparing observed metrics to their bootstrapped distributions. Multiple testing correction was performed using the Benjamini-Hochberg procedure:

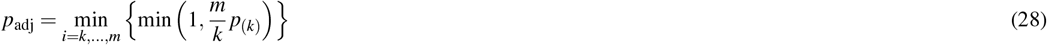

where *p*_(*k*)_ represents the *k*th smallest p-value among *m* tests.

This analysis framework was implemented using the `spatstat` package for point pattern analysis and `sf` for spatial data handling. For computational efficiency, bootstrap analyses were parallelized using the `parallel` and `doParallel` packages in R. Regions containing fewer than three cells of a given subtype were excluded from the analysis to ensure reliable statistical measurements.

### Differential Gene Expression Analysis

Visual scene group DEG was assessed via ANOVA analysis with Bonferroni correction for multiple comparisons. This analysis was conducted in Igor Pro, and involved computing the mean gene expression values of genes that passed a minimum gene expression threshold. The ANOVA compared each of the four groups against each other for a gene of interest.

Differential gene expression analysis was performed using the Model-based Analysis of Single-cell Transcriptomics (MAST) method^75^ as implemented in the `Seurat` package. For each mask-defined comparison, we constructed a `Seurat` object from the normalized expression matrix of 300 assessed genes. Data were log-normalized and scaled using default parameters. Differential expression testing was conducted using the `FindMarkers` function with MAST, setting a minimum gene detection threshold of 1% (min.pct = 0.01) across cells. For each gene *g*, we computed the log_2_ fold change (log_2_ FC) between mask-positive and mask-negative populations, along with raw and adjusted *p*-values.

Genes were considered significantly differentially expressed if they met dual criteria of |log_2_ FC| > 0.26 and adjusted *p*-value < 10^−10^. This analysis was performed both within individual RGC subtypes and across the pooled RGC population. For each mask comparison, we calculated both the percentage of cells expressing each gene within (pct_*M*_) and outside 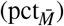 the mask region, as well as their respective mean expression values (*µ*_*M*_ and 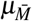).

### Primate Regression

Regional classification of the retinal surface was performed using density-based hierarchical clustering. For each pixel *p* in the normalized density grid, we constructed a feature vector **d**_*p*_ containing the normalized density values across all 45 RGC subtypes:

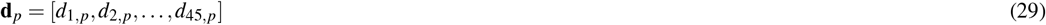

where *d*_*i*,*p*_ represents the normalized density of subtype *i* at pixel *p*. Following our previously described ensemble clustering approach, we performed hierarchical clustering using all combinations of distance metrics *D* ∈ {Euclidean, Manhattan, Maximum, Canberra, Binary, Minkowski} and linkage methods *L* ∈ {complete, average, single, Ward.D2}. For each combination of *D* and *L*, we computed cluster assignments across a range of *k* values. The optimal number of clusters *k*^∗^ was determined using silhouette analysis as before.

Human and macaque retinal single-cell RNA sequencing datasets were filtered to genes with verified mouse orthologs using biomart mapping. For each species, cells were grouped by spatial location (fovea/area centralis vs periphery). We computed log_2_ fold changes (LFC) between spatial regions for each gene *g*:

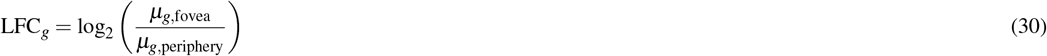

where *µ*_*g*,region_ represents the mean expression of gene *g* in the specified region. Cross-species comparisons were performed by regressing spatial LFC values between species pairs. The strength of cross-species relationships was quantified using both the coefficient of determination (*R*^2^) and Spearman’s rank correlation coefficient (*ρ*). For mouse data, the ART region that is hypothetically homologous to the area centralis identified through our hierarchical clustering approach was used to define the specialized central region analogous to fovea/area centralis in primates. The same LFC calculations and cross-species correlation analyses were performed using this computationally defined region.

### Computational Resources

All R and Python based analysis were conducted locally using consumer-grade components including an Intel i7-12700KF CPU, an NVIDIA RTX 3060 GPU with 12 GB of vRAM, and 128 GB of RAM (comprising four 32 GB Corsair CMK64GX5M2B5200C40 modules), all mounted on an MSI Z690-A Pro motherboard. The system operated on Ubuntu 22.04. All NNs and experiments were executed locally within a Python v3.9.16 Conda environment, utilizing `Jupyter Notebook` v6.5.4 as an IDE or bash commands. The primary codebase was developed using `NumPy`, `pandas`, `pyreadr`, `PyTorch`, `scikit-learn`, `tifffile`, and `tqdm`. Further data analysis and visualization were carried out in R 4.3.3, using the `RStudio` 2023.03.0 IDE with the following libraries: `clusterSim`, `cluster`, `concaveman`, `cowplot`, `deldir`, `dendextend`, `doParallel`, `dplyr`, `foreach`, `ggdendro`, `ggplot2`, `ggpubr`, `ggrepel`, `grid`, `gridExtra`, `patchwork`, `proxy`, `readr`, `Seurat`, `sf`, `smacpod`, `smerc`, `spatstat`, `spdep`, `tiff`, `tidyr`, `tidyverse`, `umap`, and `viridisLite`. Finally, Igor Pro 9 (Wavemetrics, www.wavemetrics.com) PC on Windows 11 was used for main figure preparation and analysis as indicated.

## Data and Code Availability

All custom analysis code developed for this study has been deposited in GitHub (github.com/sbudoff/MappingAllMurineRGCs). The spatial transcriptomics data generated using the 10x Xenium platform has been deposited in [database accession ID pending, please contact us prior to reviewed publication].

Single-cell RNA sequencing data from previous studies were accessed through the Single Cell Portal (singlecell.broadinstitute.org) under accession numbers SCP839, SCP509, SCP2559, and SCP919. Human retinal scRNAseq data were accessed through the CELLxGENE database (cellxgene.cziscience.com/e/d319af7f-be2e-441e-8caa-3b8a88480e89.cxg/), which extends the Single Cell Portal project SCP212.

Gene orthology relationships were established using the ENSEMBL BioMart tool (accessed through may2024.archive.ensembl.org/biomart/martview/) with manual curation and validation via the HUGO Gene Nomenclature Committee database (www.genenames.org).

## Acknowledgments

We thank the Human Immune Monitoring Shared Resource (RRID:SCR_021985) within the University of Colorado Cancer Center (P30CA046934), and in particular Mark Butters, for their expert assistance with the 10X Genomics Xenium instrument. This work was supported by the National Institutes of Health (R01EY030841, R01EY035293)

## Author contributions statement

Both authors conceived of the experiments, analyzed the results, wrote and reviewed the manuscript. S.B. conducted all wet lab work. A.P. secured all funding.

## Additional information

