## Supplemental Figures for "A Complete Spatial Map of Mouse Retinal Ganglion Cells Reveals Density and Gene Expression Specializations"

### Supplemental Registration Validation

To further validate our IHC and Xenium registration, we leveraged the vasculature stain TL and the endothelial or pericyte markers chosen by GraSP, including Pdgfrb and Cdh5. We found near perfect alignment of these endothelial and pericyte markers with the TL signal (Figure S1). Using the study regions assessed for local statistics (Figure S7), we examined the relationship between RGCs and vasculature by two methods. First, we manually annotated vascular centroid lines in Igor Pro and computed the nearest neighbor distance from each point on this line to surround RGC subtypes. With an ANOVA, we failed to reject the null hypothesis of statistically significant differences in the density of one subtype vs another within a distance of 5 $\mu$m. We next computed the count of each RGC subbtype within this distance to the vasculature and performed a bootstrap label permutation for each subtype to establish a null distribution as we did for the other local statistical assessments. Doing this revealed that only T33 demonstrated significantly greater counts density proximal to vasculature than other subtypes (Figure S7).

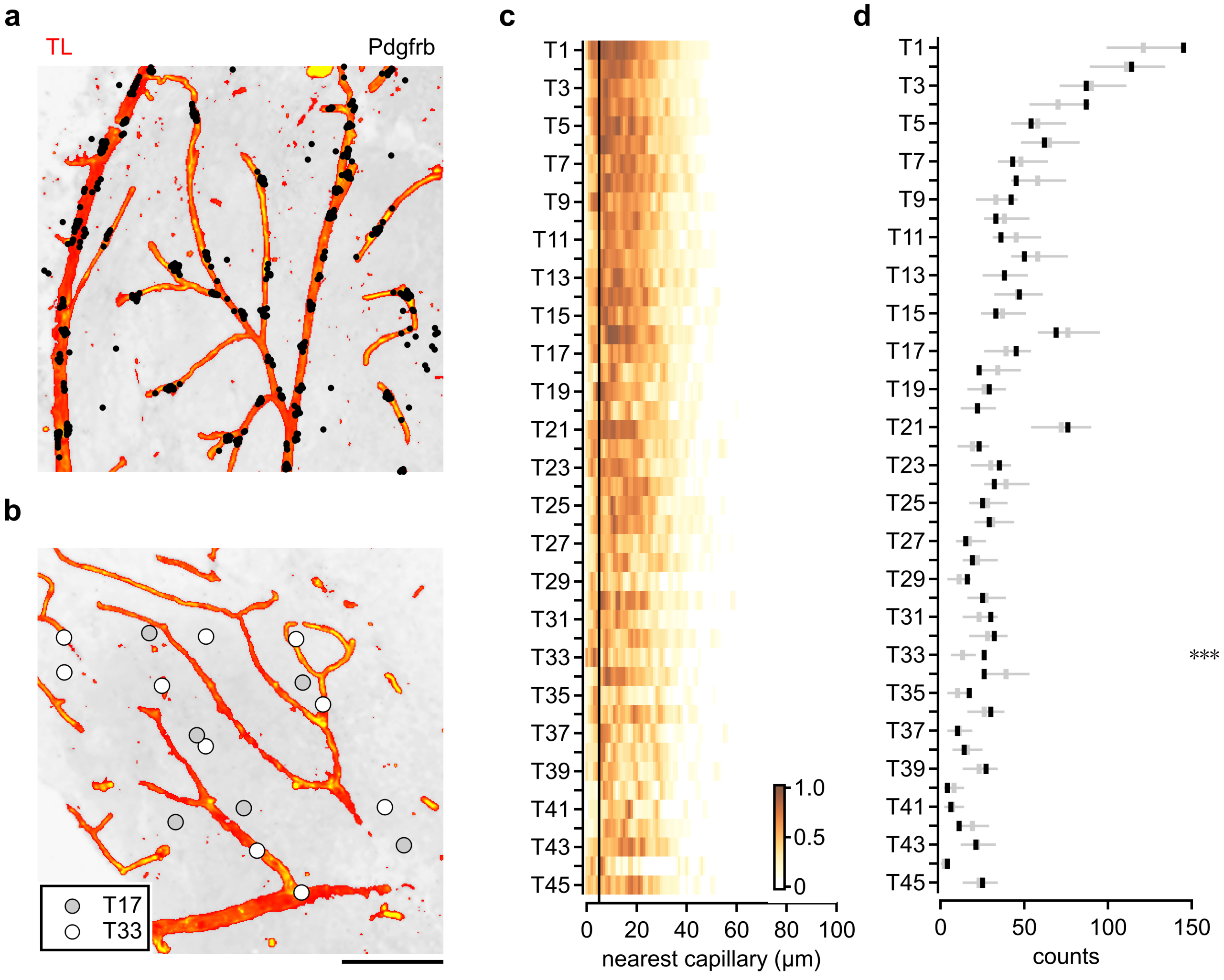


**Figure S1. Alignment of Xenium data and vasculature. a.** Overlay of tomato lectin (TL) stain that makes the vasculature with Xenium puncta for the vasculature marker platelet-derived growth factor receptor beta (Pdgfbr) showing near-perfect registration. **b.** Vasculature IHC relative to positions of T33 and T17 subtypes. Scale bar – 100 $\mu$m. **c.** Distance of each RGC subtype to the nearest capillary, calculated from a manual annotation of TL signal. **d.** Number of cells of a given subtype within 5 $\mu$m of the nearest capillary. Grey indicates bootstrap sampling derived empirical null distribution. Statistical analysis with t-test and Bonferroni correction for multiple comparisons, *** $p < .001$

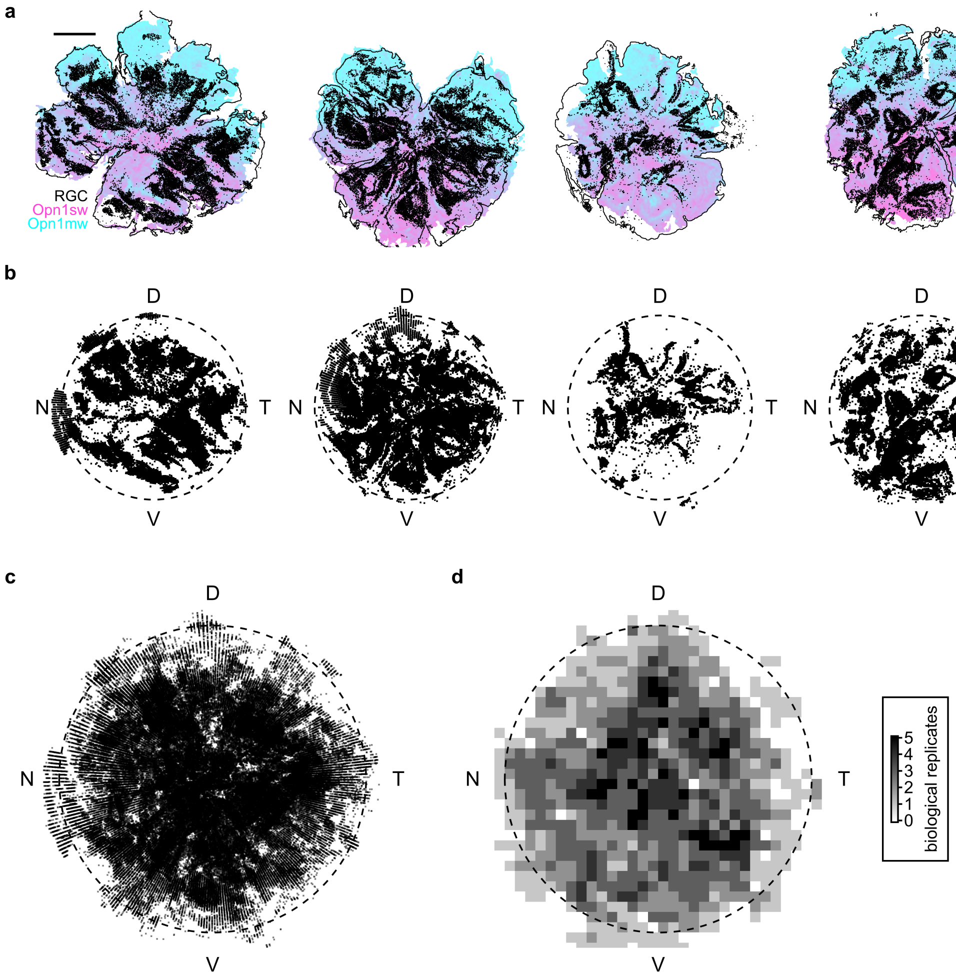


**Figure S2. Biological replicates and retinal coverage. a.** Spatial distribution of RGCs (black) overlaid over cone markers (Opn1sw for s-cone and Opn1mw for m-cone) in five examined retinas. **b.** RGC distribution for the examined retinas, corrected for relief cuts and placed in a common reference frame. The dotted circle shows an estimate of 90% of the retinal expanse. **c.** Superimposed data from **b. d.** The number of biological replicates for all locations, computed as more than 5 cells in 40 $\mu$m x 40 $\mu$m square regions.

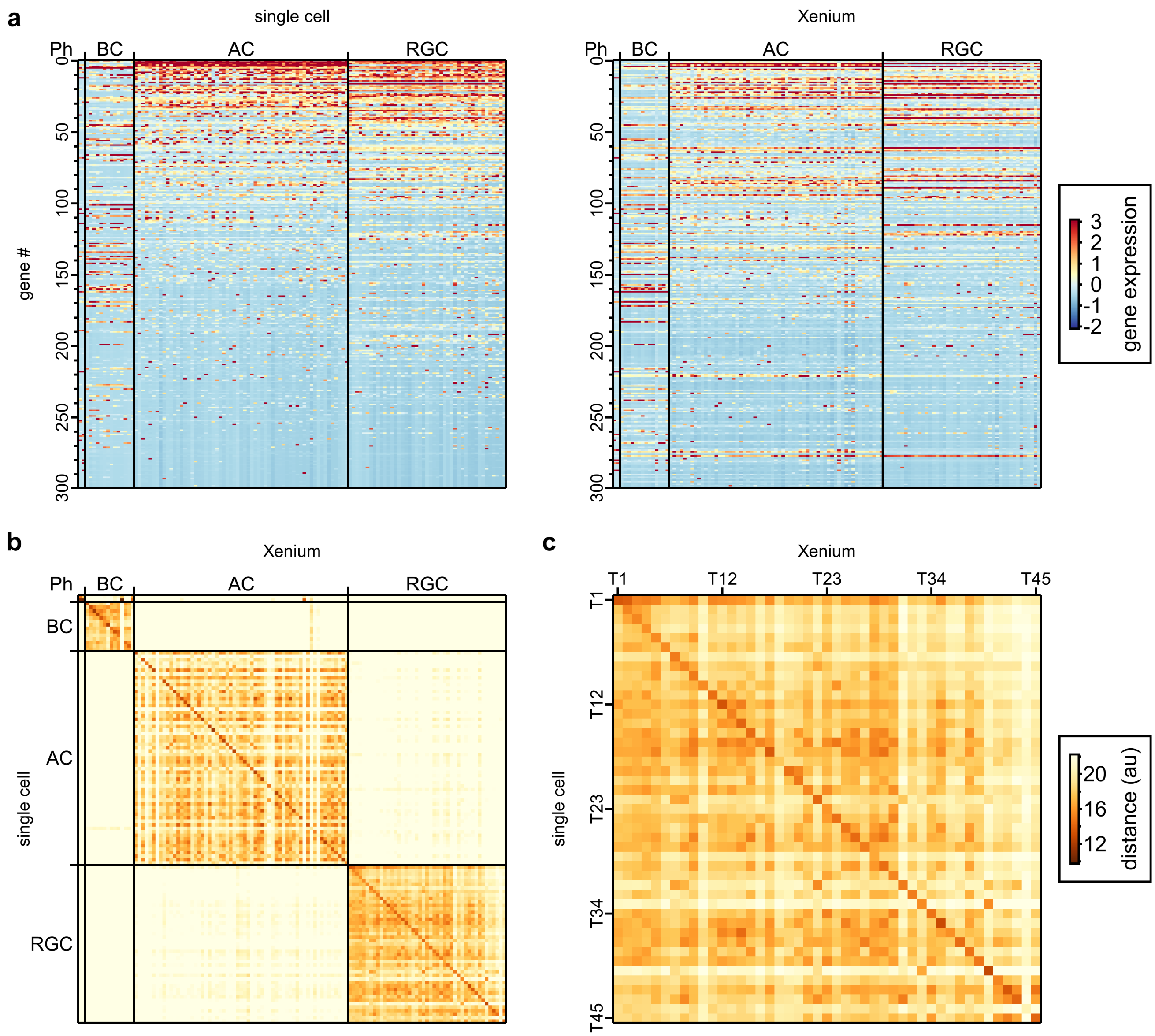


**Figure S3. Retinal cell discrimination from the 300 gene panel used in this study. a.** Left, Z-scored expression of GraSP selected gene panel of genes measured from scRNAseq datasets. Right, Z-scored expression of GraSP selected gene panel of genes measured from Xenium after Baysor segmentation. **b.** Cosine distance score of 300 gene vector between each subtype in scRNAseq dataset and Xenium. **c.** As in **b**, a zoomed-in view on RGC subtypes.

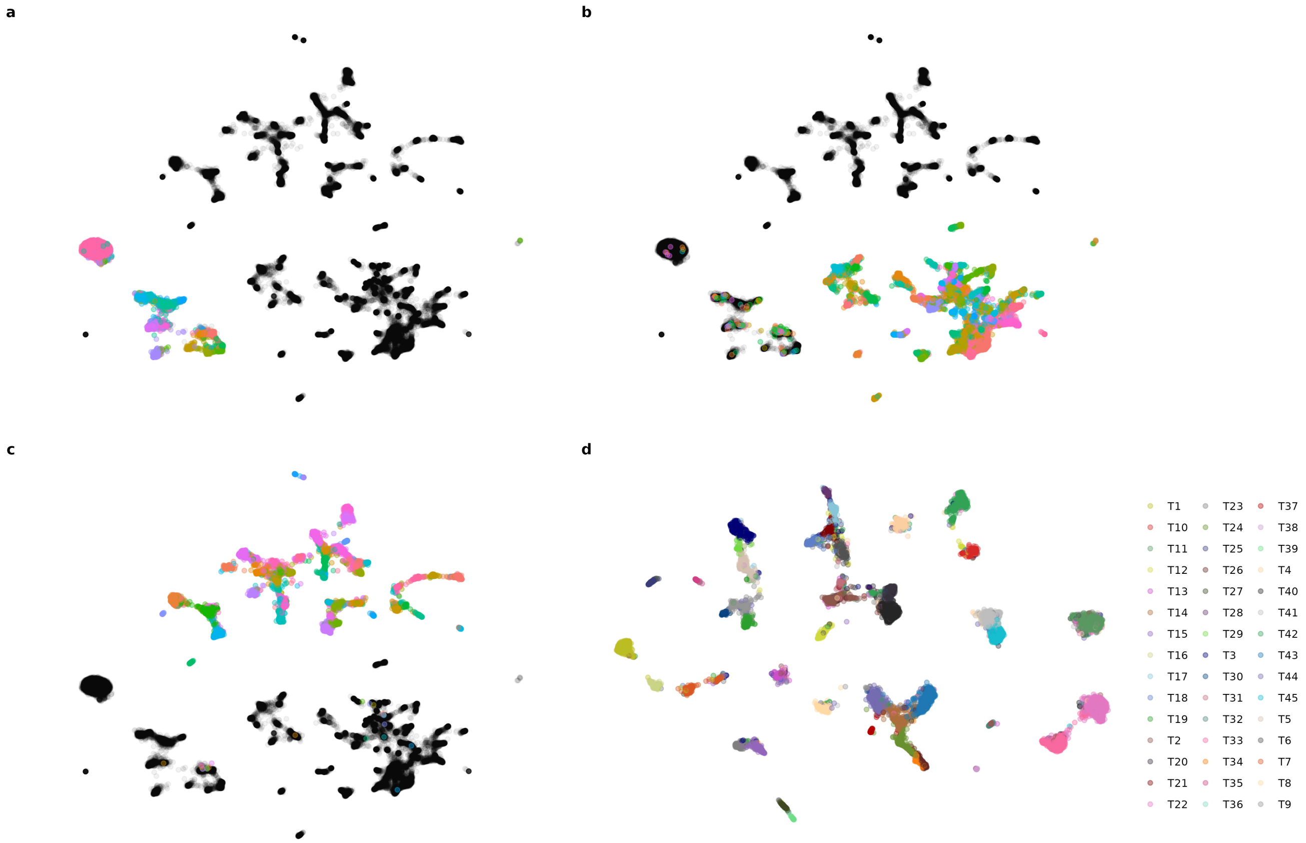


**Figure S4. UMAP projection of retinal cell subtype classification by CuttleNet. a-c.** Projection of UMAP space for the 225 genes chosen by GraSP in all retinal cells, each panel colored by one cell class based on CuttleNet inference (**a**, bipolar cells, **b**, amacrine cells, **c**, RGCs). **d.** UMAP projection of RGCs.

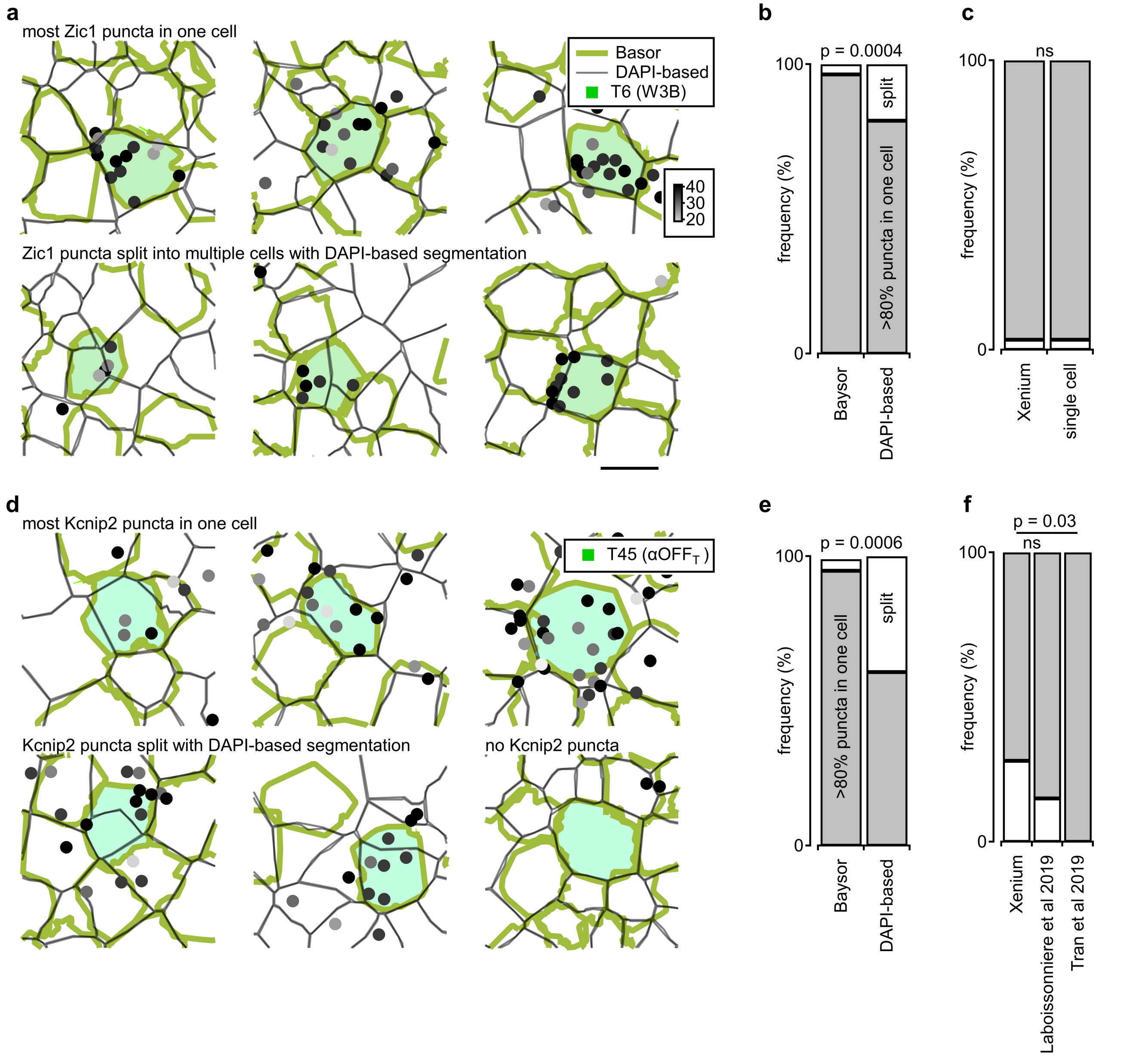


**Figure S5. Examination of RGC segmentation accuracy. a.** Distribution of Zic1 puncta, a specific and selective marker for T6 (W3B) RGCs, overlaid on segmentation results. Zic1 puncta are color-coded based on Xenium quality control metrics. Green and black curves represent Baysor and Xenium's built-in DAPI-based segmentations, respectively. Inferred T6 RGCs are shaded in green. The top row shows examples where Zic1 puncta clusters are correctly assigned to a single cell in both segmentations. The bottom row highlights cases where Zic1 clusters are divided among multiple cells by the Xenium algorithm but assigned to a single cell by Baysor segmentation. **b.** Frequency distribution of Zic1 puncta clusters assigned to a single cell (gray) or split among multiple neighboring cells (white). The analysis is based on 87 manually curated RGCs. Statistical significance was assessed using a chi-square test. **c.** Comparison of the fraction of W3B cells expressing Zic1 between this Xenium dataset (gray) and single-cell data, showing consistent expression levels. **d-f.** As in **a-c.** for Kcnip2, which is a less-specific, but commonly used marker of T45 ($\alpha$OFFT) RGCs[25,41,46]

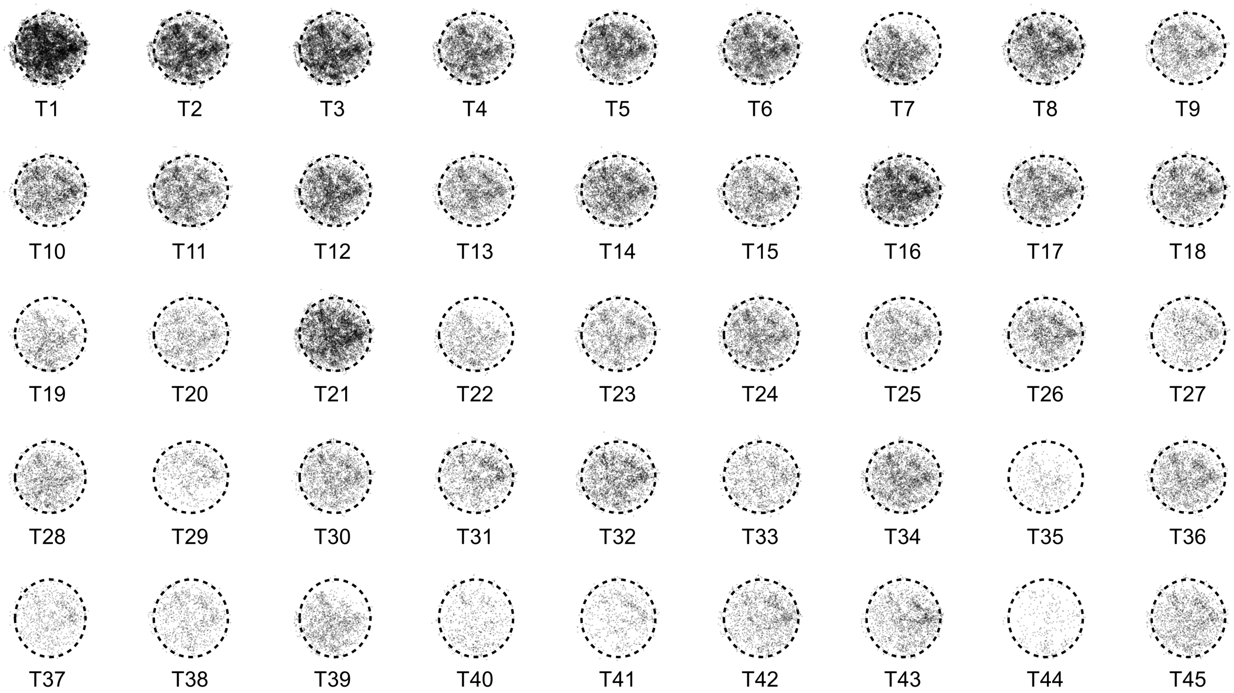


**Figure S6. Point process maps of all RGC subtypes in the circularized reconstructed retinas.** The dashed line represents the study region as in Figure S2

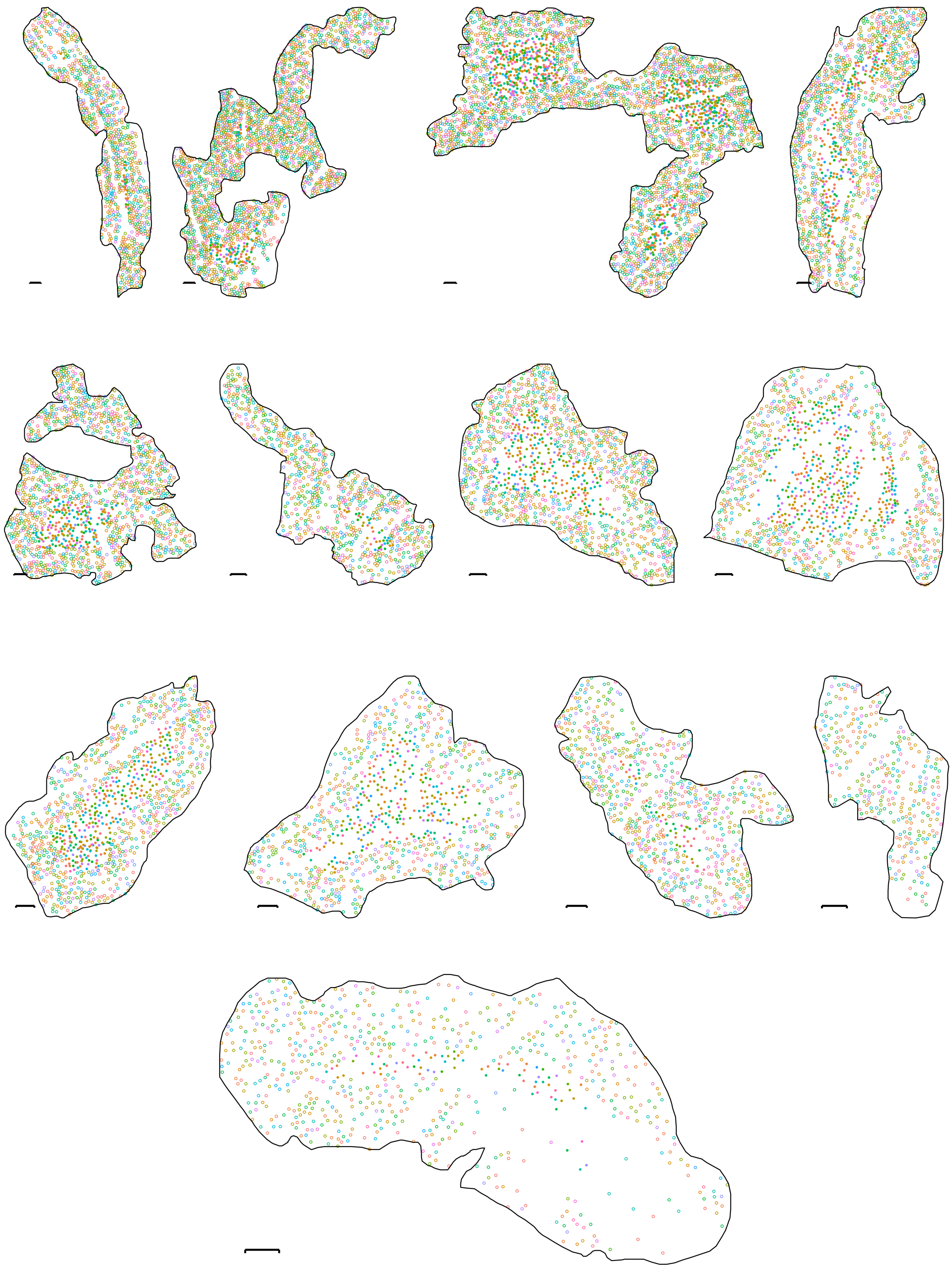

**Figure S7. 14 Manually Selected Study Regions for Local Statistical Analysis.** Scale bars – 50 $\mu$m. Points colored by subtype classification, as in Figure S4. Points with solid fill were determined by `spatstat` to be within the study region, cells outside this algorithmically defined boundary are marked with opaque circles
.

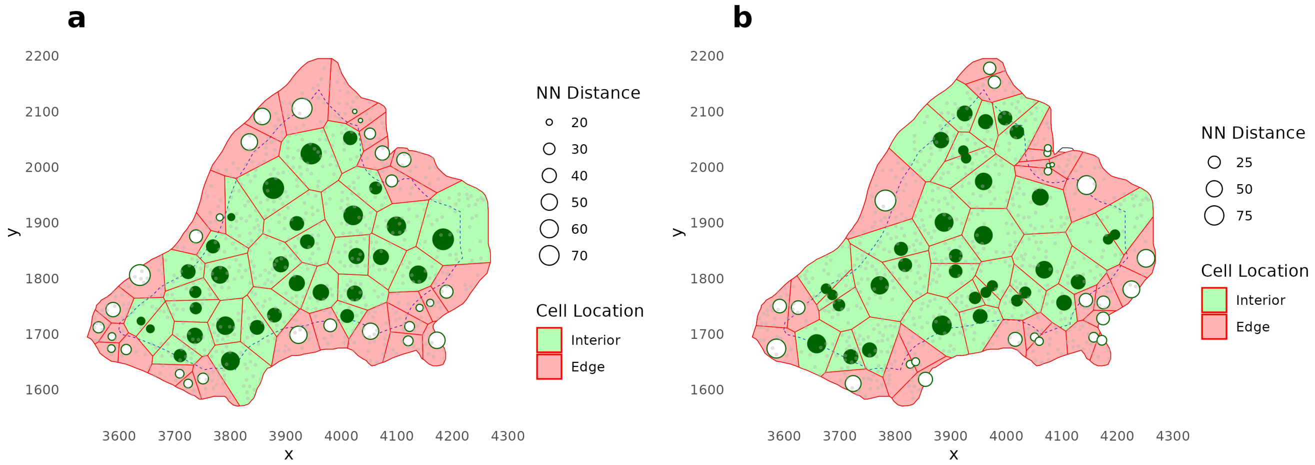


**Figure S8. Demonstration of local statistic computations. a.** The example illustrates the analysis of T1 RGCs (circles, with size scaled by the nearest neighbor distance) in a representative study region. Solid lines indicate Voronoi tessellations. Study region edge from `spatstat` (dashed lines, red), are excluded from analysis. **b.** as in **a**, but for bootstrap sampling example of local statistic calculations for an equal number of cells drawn without replacement from the full RGC population (grey dots).

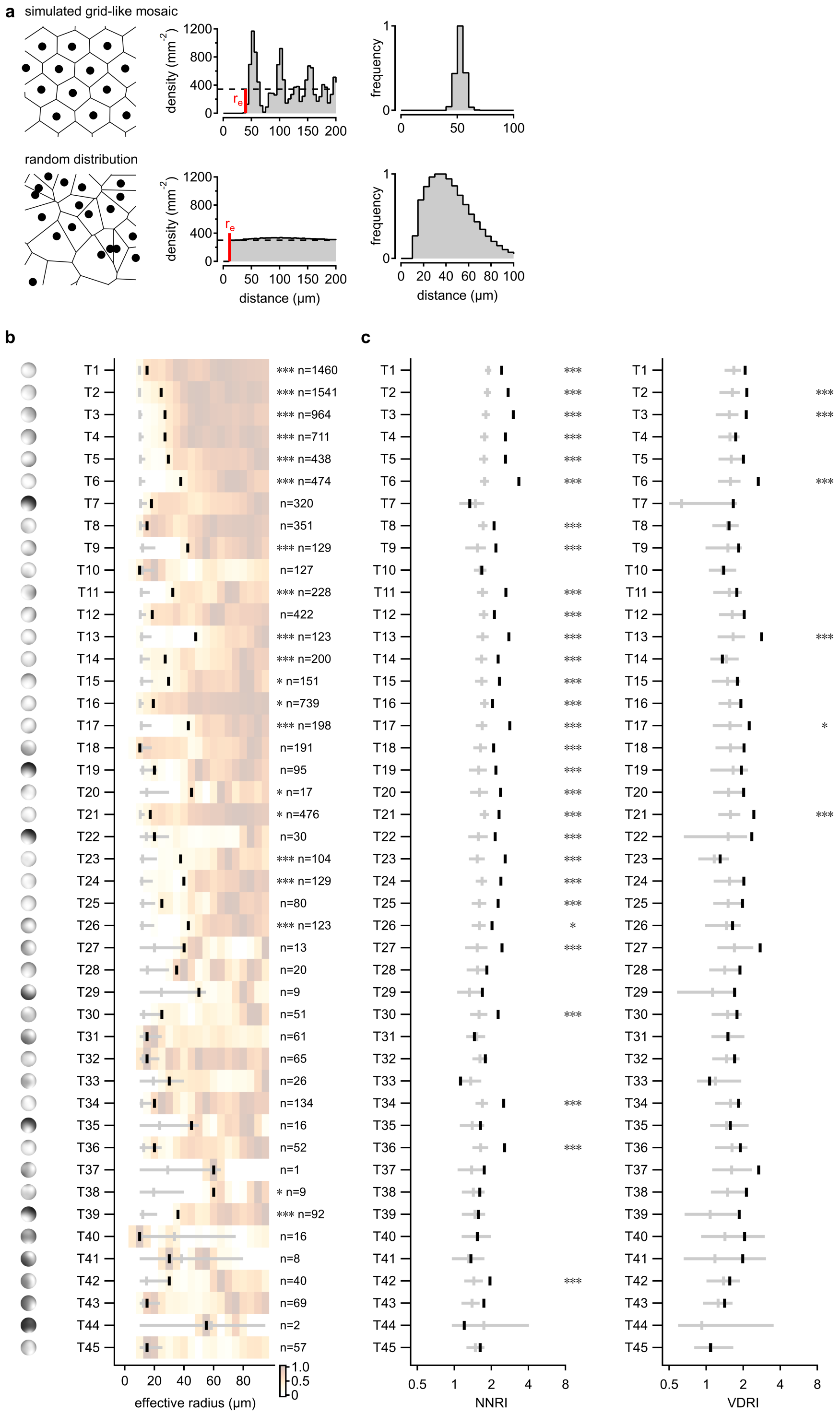

**Figure S9. Local statistics of all RGC subtypes. a.** Top, simulation of a perfectly packed, hexagonal lattice showing the autocorrelation density plot and $r_e$ as well as the distribution of nearest neighbor distances. Bottom, simulation of a completely spatially random set of points with the same distribution shown. **b, c.** as in Figure 3e-g with all 45 cell subtypes shown. $n$, the number of cells in all study regions. $^{*}p < .05$, $^{***}p < .001$.

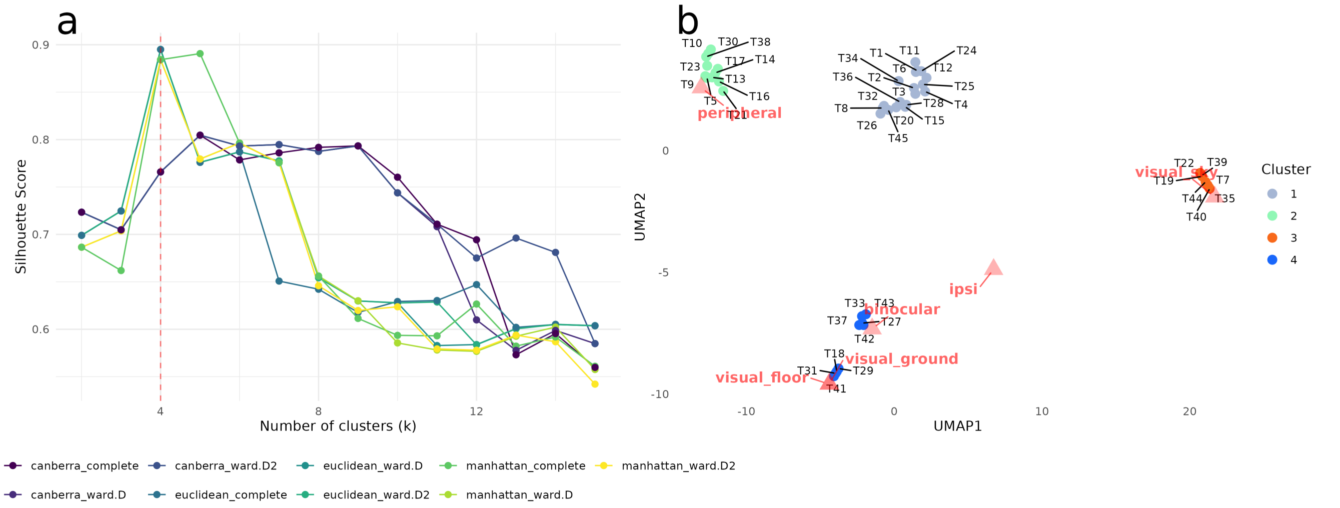


**Figure S10.** **Unsupervised selection of RGC subtype clustering. a.** Silhouette analysis for hierarchical clustering conducted with 3 different distance calculations and 3 different linkage methods. $k^* = 4$ was found as the optimal cluster number. **b.** UMAP projection of $F_1$ score vectors for all RGCs with cluster assignments by ensemble vote from hierarchical clustering algorithms. Color coding as in Figure 4. To visualize the manifold's practical implications, the visual scene masks' $F_1$ scores were also computed in the same manner and thier vector representation was co-projected, masks position on the manifold is labeled with red triangles.

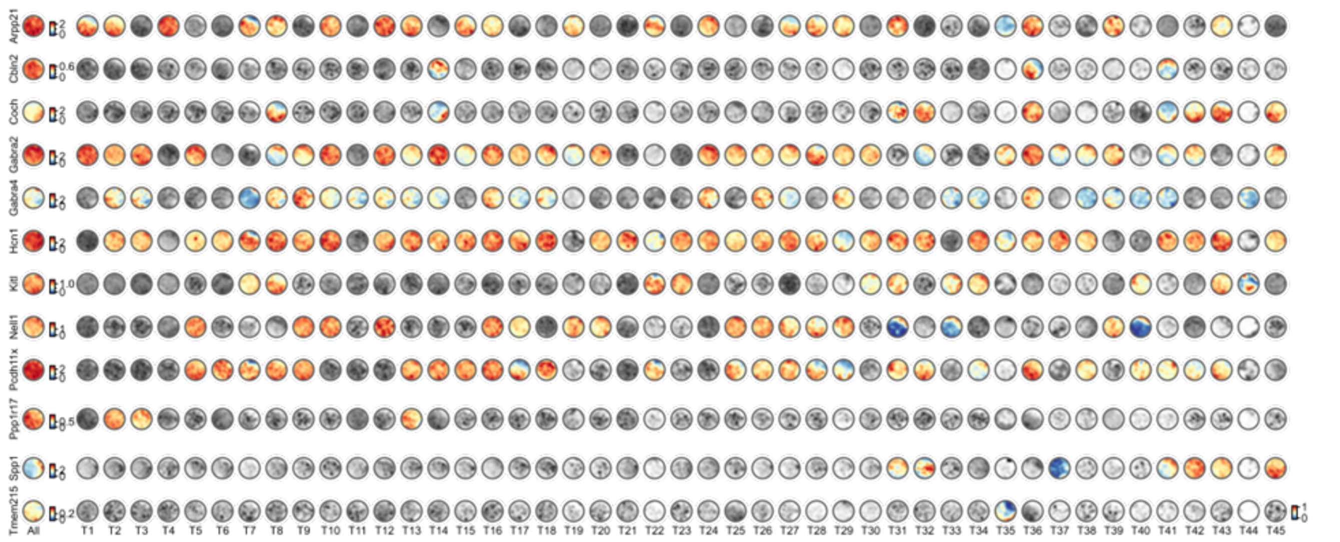


**Figure S11.** **Representative topography of spatially variable DEGs in all RGC subtypes.** Maps of genes with <1 puncta per cell for the given subtype are colored in greyscale.

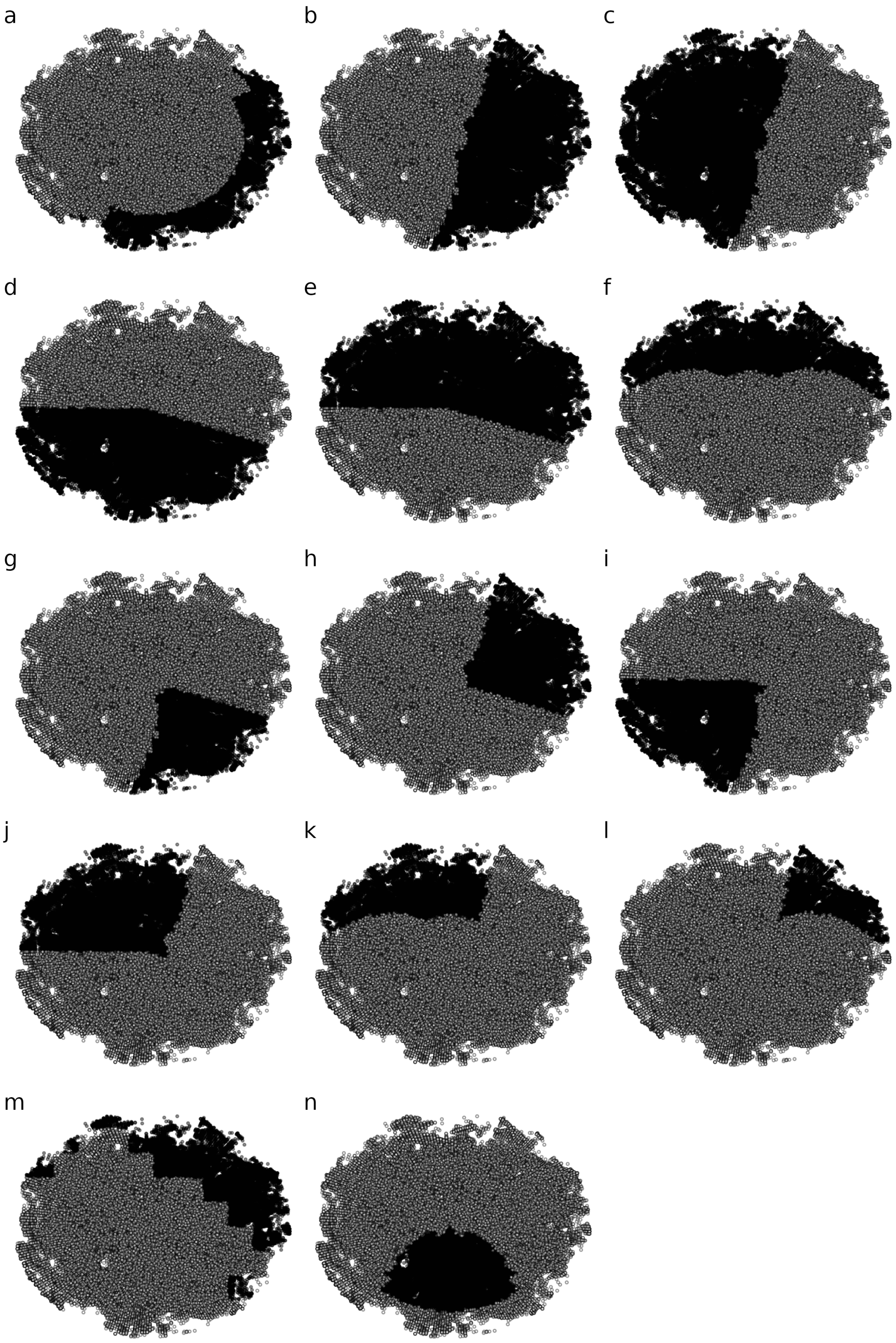


**Figure S12. Visual scene masks.** Visual scene masks derived from: **a.** ipsilateral projection RGCs, **b.** binocular, **c.** peripheral, **d.** sky, **e.** ground, **f.** floor, **g.** binocular ∪ sky, **h.** binocular ∪ ground, **i.** peripheral ∪ sky, **j.** peripheral ∪ ground, **k.** peripheral ∪ floor, and **l.** binocular ∪ floor regions. **m.** Mask for the *area retinae temporalis* (ART) computed from hierarchical clustering of RGC subtype density vector. **n.** Mask for the *area centralis* estimated from RGC density maps[24], with a similar area as the mask in **m**.

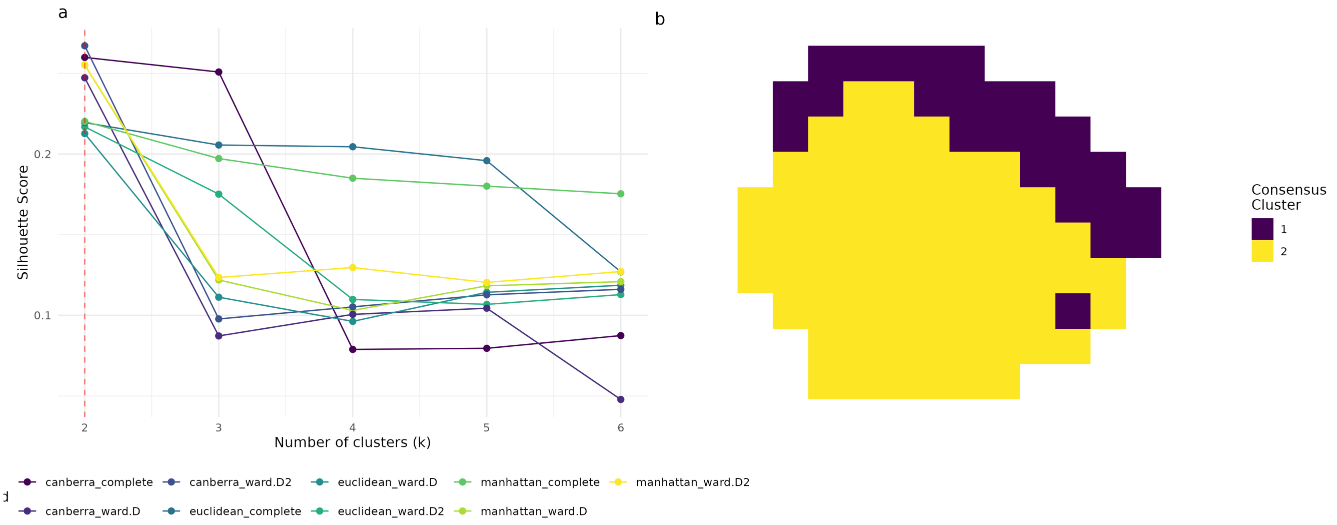


**Figure S13. Hierarchical clustering of subtype proportionality across the retina converges on an ART-like region being different from the rest of the retina. a.** Silhouette analysis of an ensemble of hierarchical clustering models using different distance and linkage methods. Optimal clustering was seen with $k^* = 2$ **b.** Consensus clustering projected onto retinal coordinates.

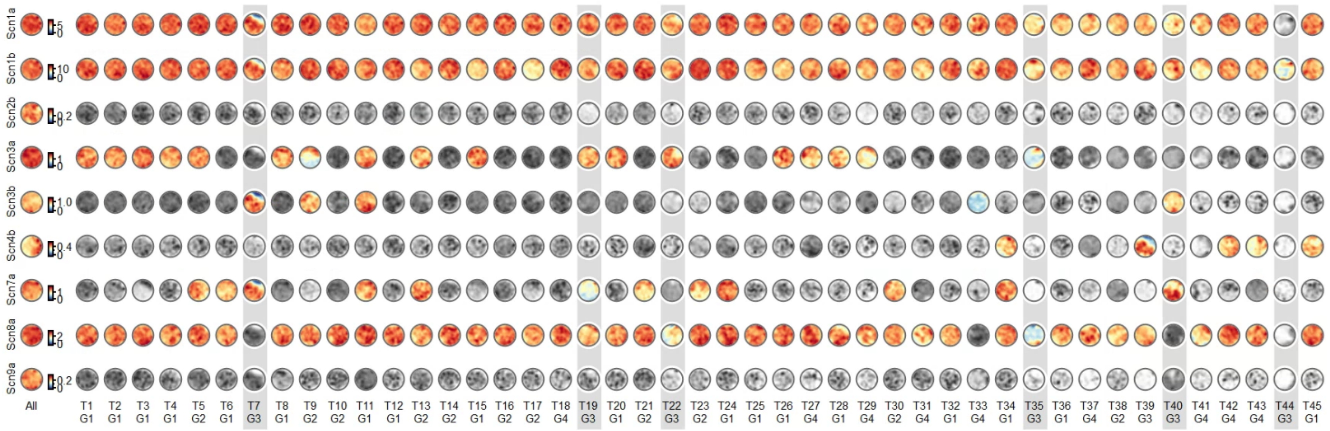


**Figure S14. Spatial distribution of voltage gated sodium channels suggests correlation between primates and mice driven by group 3 subtypes.** Gene topography across all RGCs. Group 3 subtypes highlighted in grey background. Maps of genes with <1 puncta per cell for the given subtype are colored in greyscale.

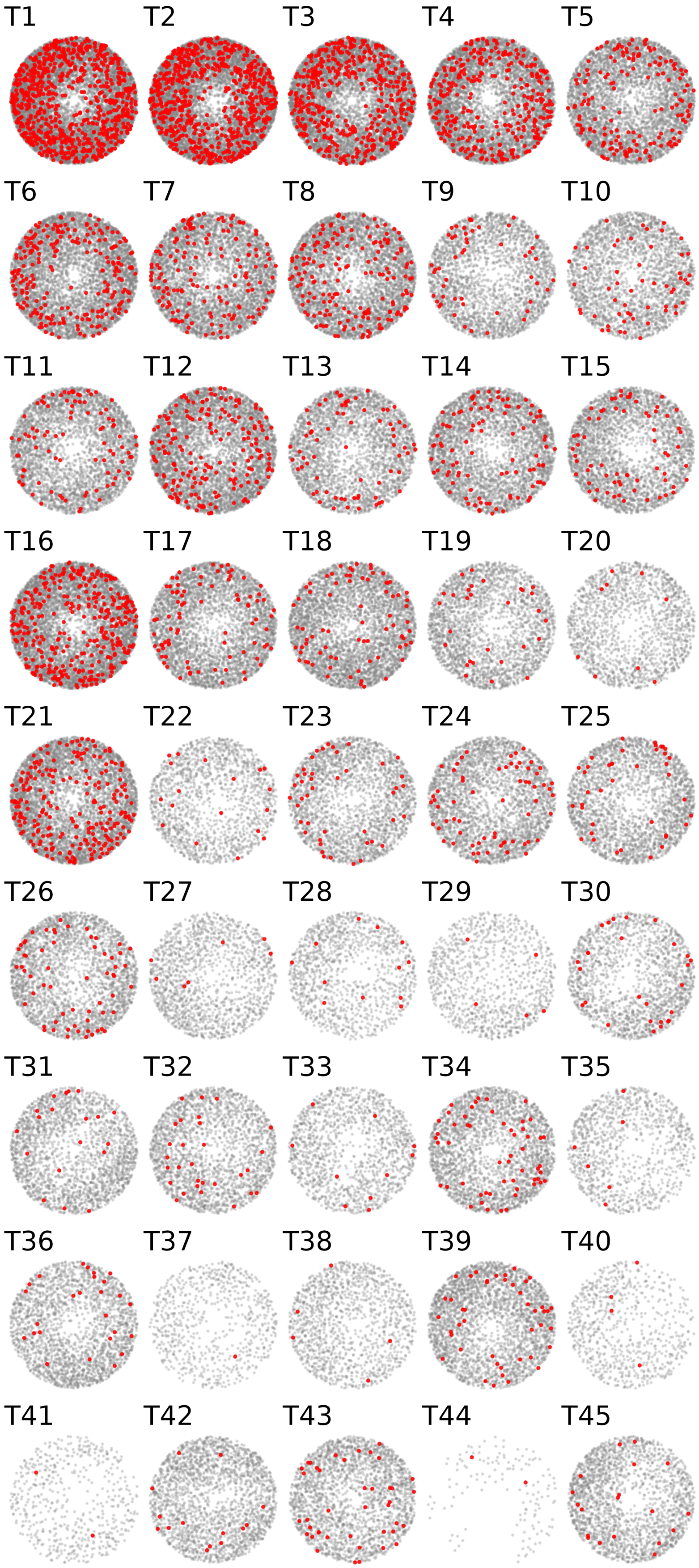


**Figure S15.** **Autocorrelation of each subtype (Red) shown against against all other RGCs (grey).**
